# Temporal analysis of physiological phenotypes identifies novel metabolic and genetic underpinnings of senescence in maize

**DOI:** 10.1101/2025.03.07.641920

**Authors:** Manwinder S. Brar, Rohit Kumar, Bharath Kunduru, Christopher S. McMahan, Nishanth Tharayil, Rajandeep S. Sekhon

## Abstract

Leaf senescence induces extensive metabolome reprogramming to optimize nutrient recycling, enhance resilience to abiotic and biotic stress, and improve productivity. However, the characterization of these metabolic shifts and the identification of key metabolites and pathways remains limited. We generated a temporal map of physiological and metabolic diversity in genetically diverse maize inbred lines varying for the staygreen trait. Combinatorial analysis of physiological and metabolic changes revealed substantial metabolic perturbations and identified 84 leaf metabolites associated with senescence. Non-staygreen inbred lines exhibited higher accumulation of primary metabolites including sugar alcohols such as mannitol and erythritol, and amino acids such as phenylalanine and arginine. In contrast, the staygreen inbred lines showed higher abundance of secondary metabolites, primarily phenylpropanoids, including caffeic acid, chlorogenic acid, and eriodictyol. Linking metabolome to the genome identified 56 novel candidate genes expressed in adult maize leaf that regulate metabolic flux during senescence. Reverse genetic analysis validated the role of naringenin chalcone and eriodictyol in both maize and Arabidopsis, demonstrating a conserved function of these phenylpropanoids in leaf senescence across monocots and dicots. Our study provides valuable insights into the coordinated physiological and metabolic changes driving leaf senescence and identifies novel genes underlying this complex developmental process.

## INTRODUCTION

Senescence is an age-dependent process that leads to the death of a cell, tissue, or organ and, in the case of annual plants, culminates in the death of the whole plant. Leaf senescence is the final phase of leaf development wherein programmed events lead to the degradation of cellular macromolecules and the remobilization of nutrients to developing tissues or storage organs (Lim et al., 2007). Timely senescence ensures the efficient recycling of leaf nutrients, particularly nitrogen (N), to developing organs, including grains and contributes to grain quality and improves overall plant fitness (Buchanan-Wollaston, 1997; Mayta et al., 2019; Guo et al., 2021). However, leaf senescence compromises photosynthetic activity, limits the duration of the carbon-capture phase, and results in a loss of net carbon productivity (Noodén, 1988). Delayed leaf senescence, a trait referred to as staygreen (or stay-green), is associated with greater leaf longevity, prolonged photosynthesis, higher grain yield, and increased total dry matter (Lee and Tollenaar, 2007; Duvick et al., 2010; Gregersen et al., 2013; Thomas and Ougham, 2014; Wang et al., 2023). The staygreen crop plants exhibit enhanced drought and heat tolerance and increased resistance to diseases, especially those caused by stalk-rotting pathogens (Bertolini et al., 1983; Ougham et al., 2007; Vijayalakshmi et al., 2010; Jordan et al., 2012; Emebiri, 2013). Given that agricultural production systems are increasingly challenged by the rapidly worsening climate and associated extreme weather events, the staygreen trait is becoming increasingly valuable for sustainable crop production. While the biological and agronomic significance of senescence is clear, the mechanisms regulating the onset and progression of this complex process in maize (*Zea mays* L.) and related grasses remain poorly understood.

While the visual manifestation of the staygreen trait is similar across species and genotypes, characterized by delayed senescence and prolonged greenness, the underlying biological mechanisms and genetic determinants are diverse (Thomas and Howarth, 2000). Therefore, a comprehensive mechanistic understanding of staygreen in a species relies on differentiating the physiological and metabolic processes underlying the mechanistic diversity of the trait. Senescence is a biologically active process initiated and accelerated by continuous integration of internal and external signals and manifested as a spatial and temporal continuum of physiological phenotypes (Lim et al., 2007; Thomas, 2013; Bresson et al., 2018; Kumar et al., 2023). Diverse aspects of leaf physiology used as indicators of senescence include visual scoring of leaf greenness, quantification or estimation of chlorophyll content, assessment of photosystem II photochemistry, and CO_2_ assimilation (Großkinsky et al., 2017; Bresson et al., 2018). Since these physiological indicators are related to distinct cellular processes in leaf cells, the extent and nature of the staygreen trait captured by these phenotypes are expected to be diverse. Furthermore, recording these phenotypes at a given time point without considering the impact of their temporal nature and variable duration may also obscure the genetic analysis and mechanistic understanding of senescence. For instance, while a snapshot of chlorophyll content fails to capture the phenotypic variation for the staygreen trait, temporal changes in chlorophyll content and photosynthetic activity can discern genotypes ranging from functional to cosmetic staygreen (Thomas and Howarth, 2000).

Leaf metabolome undergoes a dramatic change during senescence with the transition of leaves from carbon (C) source to nitrogen (N) source to transport nitrogen to the sink tissues, primarily developing seeds (Kumar et al., 2019). Dramatic changes in C- and N-containing metabolites and their flux among sources and sinks are, therefore, at the crux of senescence. Certain metabolites also act as signaling molecules to modulate genome-wide transcription and translation and, therefore, dictate the initiation and progression of senescence (Buchanan-Wollaston et al., 2005; Vidal and Gutiérrez, 2008; Smeekens et al., 2010; Watanabe et al., 2013; Li et al., 2017). Furthermore, the metabolome is the final readout of the central dogma and, therefore, represents a reliable link between the genome and phenome and provides unique opportunities to identify genes underlying a complex phenotype. Thus, characterization of the metabolome is crucial for understanding mechanisms that initiate and maintain nutrient partitioning during senescence. Recent studies have provided valuable information on metabolic shifts during senescence and the underlying genetic components in model and crop plants (Zhang and Becker, 2015; Guo et al., 2021; Zhu et al., 2022). However, a comprehensive understanding of metabolome changes and their association with natural genetic variation in leaf senescence in maize remains obscure.

To generate a comprehensive mechanistic understanding of the physiological indicators and metabolic processes underlying senescence, we performed a detailed analysis of a set of diverse maize inbred lines exhibiting varying levels of staygreen trait. We evaluated inbred lines using multiple physiological traits to determine a standardized high-throughput phenotyping approach for senescence. We then performed a time-course analysis of targeted and untargeted metabolome on staygreen and non-staygreen inbred lines to understand the metabolic components that regulate senescence. Finally, we linked a subset of metabolites to the genes and validated two of the genes to demonstrate the value of metabolome in uncovering novel genetic determinants of senescence. This study provides a valuable roadmap to an enhanced understanding of the genetic architecture of senescence in maize and related grasses and for employing such knowledge to tailor crops for the future.

## RESULTS

### Diversity of physiological traits related to senescence

To examine the physiological and metabolic diversity underlying senescence, we identified a set of 19 diverse maize inbred lines representing diverse heterotic groups (Table 1, Supplementary Figure S1). Of these, seven and four lines have been designated as staygreen and non-staygreen in the published records, respectively, while no information is available for the rest. To evaluate the extent and nature of senescence, we recorded five physiological traits representing diverse cellular processes during the post-flowering period and classified the inbred lines for their senescence phenotype based on each trait. Principal component analyses based on chlorophyll content estimated by SPAD and the rate of CO_2_ assimilation, the two physiological traits that may provide a direct estimate of photosynthetic activity, classified these inbred lines into two major groups (Figure 1, A-B; Supplementary Dataset S1). Of these, one group (N=11) included all seven inbred lines reported to be staygreen, while the other group (N=8) included four known non-staygreen inbred lines (Table 1). Analysis based on photosystem II activity (Fv/Fm), a trait often used to assess senescence and stress in plants, also provided a similar clustering, albeit with less distinction among the two groups (Figure 1C). These analyses corroborated the previously reported data for the eleven lines and allowed us to classify the remaining inbred lines for senescence phenotype (Table 1). Remarkably, the accumulation of reactive oxygen species, a fundamental regulator of diverse cellular processes and hallmark of oxidative stress during senescence, resulted in a tight clustering of the staygreen inbred lines while a more scattered clustering of the non-staygreen lines (Figure 1D). This observation highlights tremendous variation in the generation and/or quenching of ROS during senescence in maize. Finally, leaf senescence represents the transition of leaves from a C source through photosynthetic assimilation to an N source via proteolysis of leaf proteins followed by N remobilization (Thomas and Ougham, 2014). Interestingly, clustering based on the C/N ratio failed to discern the inbred lines (Figure 1D), indicating a substantial variation for C partitioning and N remobilization during senescence that is not tightly correlated with the staygreen trait. To summarize, the inbred lines used in the study captured striking differences in physiological traits related to senescence.

**Figure 1:**
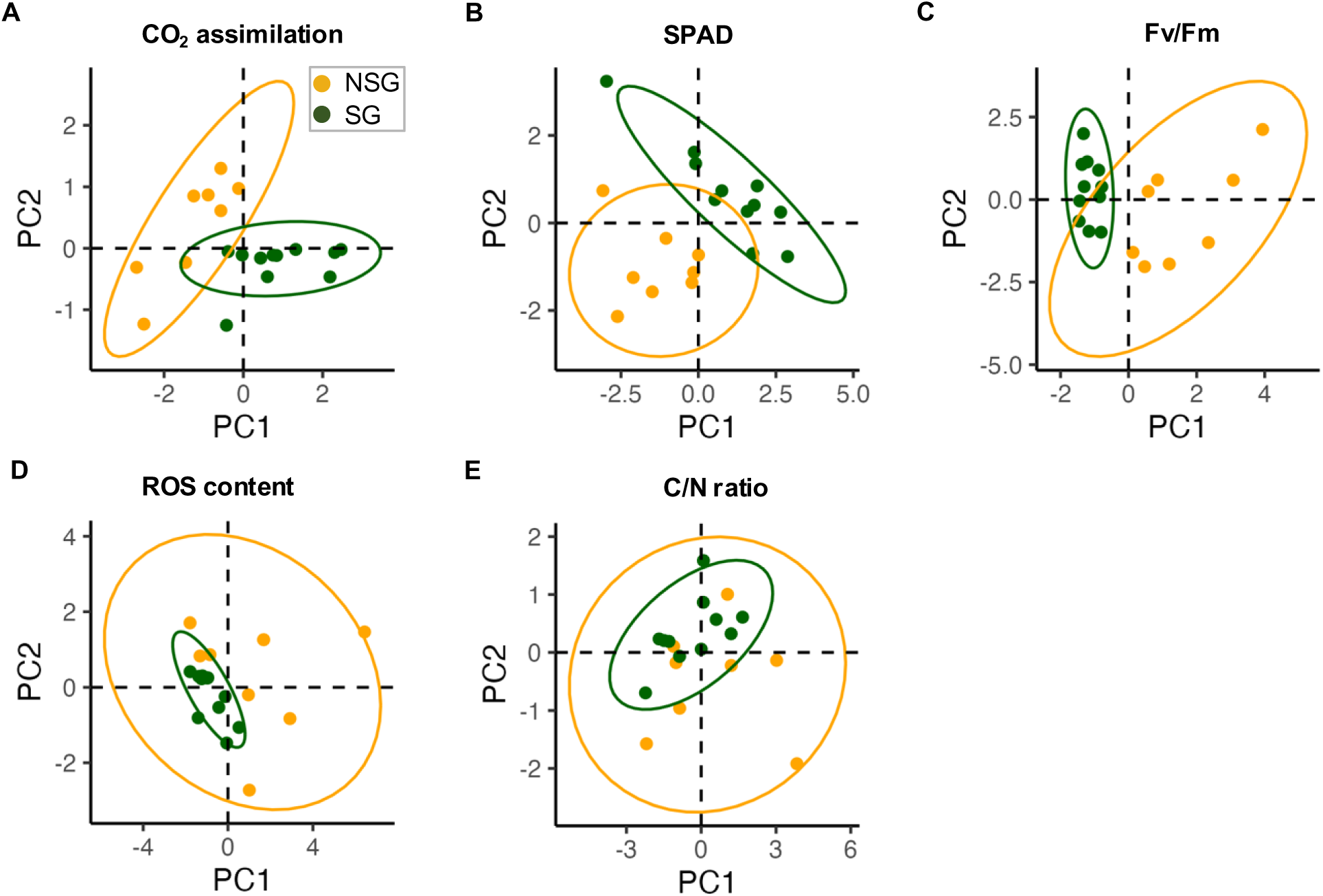
Diversity of physiological traits underlying senescence. Principal component analysis of staygreen (SG, green circles) and non-staygreen (NSG, orange circles) inbred lines based on CO_2_ assimilation (**A**), SPAD (**B**), Fv/Fm (**C**), ROS content (**D**), and C/N ratio (**E**). PC1 and PC2 denote the first and second principal components, respectively. Orange and green ovals represent the confidence interval (95%) for the non-staygreen and staygreen inbred lines, respectively.

**Table 1:**
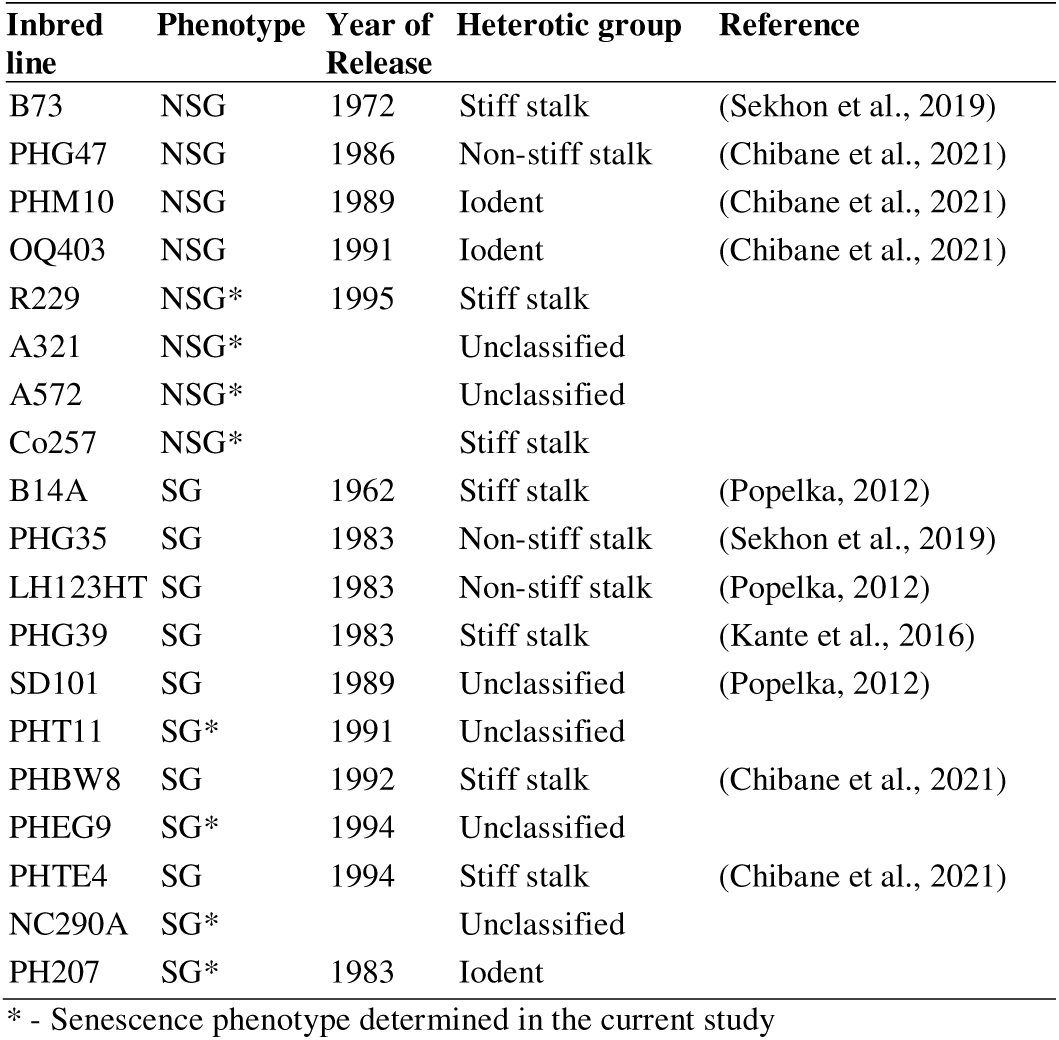
Inbred lines used in the study.

| Inbred line | Phenotype | Year of Release | Heterotic group | Reference |
| --- | --- | --- | --- | --- |
| B73 | NSG | 1972 | Stiff stalk | (Sekhon et al., 2019) |
| PHG47 | NSG | 1986 | Non-stiff stalk | (Chibane et al., 2021) |
| PHM10 | NSG | 1989 | Iodent | (Chibane et al., 2021) |
| OQ403 | NSG | 1991 | Iodent | (Chibane et al., 2021) |
| R229 | NSG* | 1995 | Stiff stalk |  |
| A321 | NSG* |  | Unclassified |  |
| A572 | NSG* |  | Unclassified |  |
| Co257 | NSG* |  | Stiff stalk |  |
| B14A | SG | 1962 | Stiff stalk | (Popelka, 2012) |
| PHG35 | SG | 1983 | Non-stiff stalk | (Sekhon et al., 2019) |
| LH123HT | SG | 1983 | Non-stiff stalk | (Popelka, 2012) |
| PHG39 | SG | 1983 | Stiff stalk | (Kante et al., 2016) |
| SD101 | SG | 1989 | Unclassified | (Popelka, 2012) |
| PHT11 | SG* | 1991 | Unclassified |  |
| PHBW8 | SG | 1992 | Stiff stalk | (Chibane et al., 2021) |
| PHEG9 | SG* | 1994 | Unclassified |  |
| PHTE4 | SG | 1994 | Stiff stalk | (Chibane et al., 2021) |
| NC290A | SG* |  | Unclassified |  |
| PH207 | SG* | 1983 | Iodent |  |

\* - Senescence phenotype determined in the current study

### Dynamic nature of physiological traits underlying senescence

To understand the timing and rate of progression of diverse physiological changes related to senescence and the association of such changes with the staygreen trait, we compared the temporal changes in three distinct physiological phenotypes. For each of the three phenotypes (SPAD, Fv/Fm, and ROS), we identified the developmental stage that showed significant change (i.e., changepoint) and estimated the rate of change for each inbred line during senescence. Remarkably, the changepoint for the three traits varied for diverse inbred lines, indicating inherent genetic variation in the underlying physiological processes manifesting these traits. For instance, chlorophyll content (SPAD) showed an earlier decline (by ∼1d) in PHG35, a staygreen inbred line, compared to a non-staygreen inbred B73 (Figure 2A). In contrast, the decrease in Fv/Fm values started earlier (by ∼2d) in B73 compared to PHG35 (Figure 2B). Interestingly, PHG35 leaves started accumulating ROS content earlier (by ∼4d) compared to B73 (Figure 2C). In contrast, the progression of various physiological changes revealed that both Fv/Fm and SPAD values declined while the ROS content increased rapidly after the changepoint in the non-staygreen inbred lines compared to the staygreen lines (Figure 2). Such a lack of association between the timing of the onset of a physiological change and staygreen trait demonstrates that relying on a snapshot of a phenotype is not an ideal strategy to reveal the inherent genetic differences. While various physiological changes related to senescence can initiate at variable times, their rate of progression is of higher significance in determining the staygreen trait. Therefore, temporal monitoring of the rate of a physiological process will provide more comprehensive insights into the genetic mechanisms underlying senescence.

**Figure 2:**
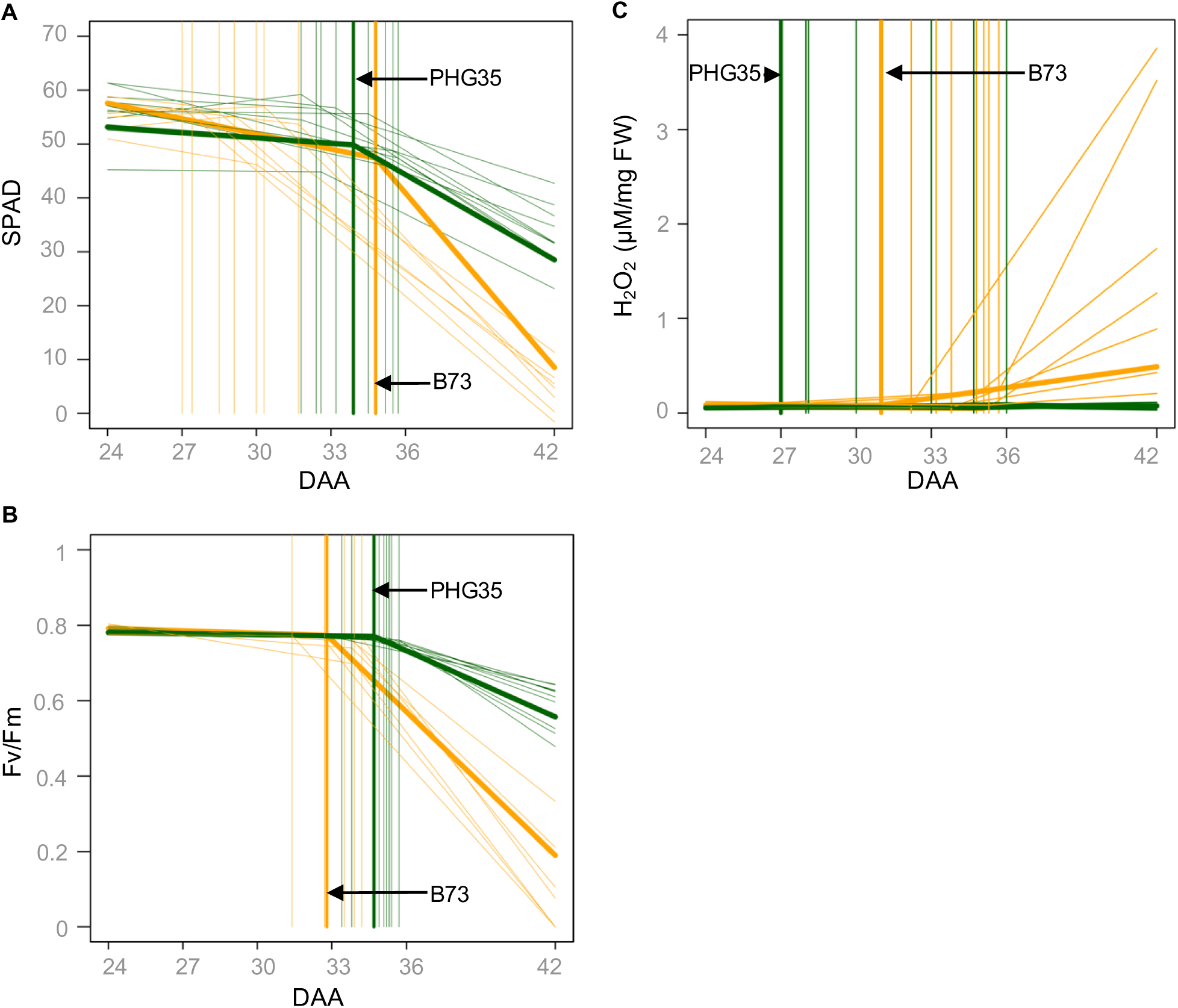
Dynamic nature of physiological traits underlying senescence. Changepoint analysis of diverse inbred lines for SPAD values (**A**), Fv/Fm (**B**), and ROS content (**C**). Vertical lines represent changepoints for different genotypes. Green and orange lines represent staygreen and non-staygreen inbred lines, respectively. DAA denotes days after anthesis.

### Temporal variation in primary metabolome underlying senescence

To capture the metabolome variation, we examined the changes in primary metabolites of five staygreen and five non-staygreen inbred lines at three leaf development stages selected based on temporal phenotypic analysis (Figure 2). The untargeted primary metabolomic analysis detected 48 metabolites belonging to four different groups (Supplementary Figure S2). Partial least square discriminant analysis (PLS-DA) based on the 48 primary metabolites did not distinguish between the staygreen and non-staygreen inbred lines, indicating a lack of global differences in primary metabolism during senescence (Figure 3A). This observation indicates that differences in senescence phenotype are not associated with global changes in primary metabolism likely due to the importance of these metabolites in plant survival.

**Figure 3:**
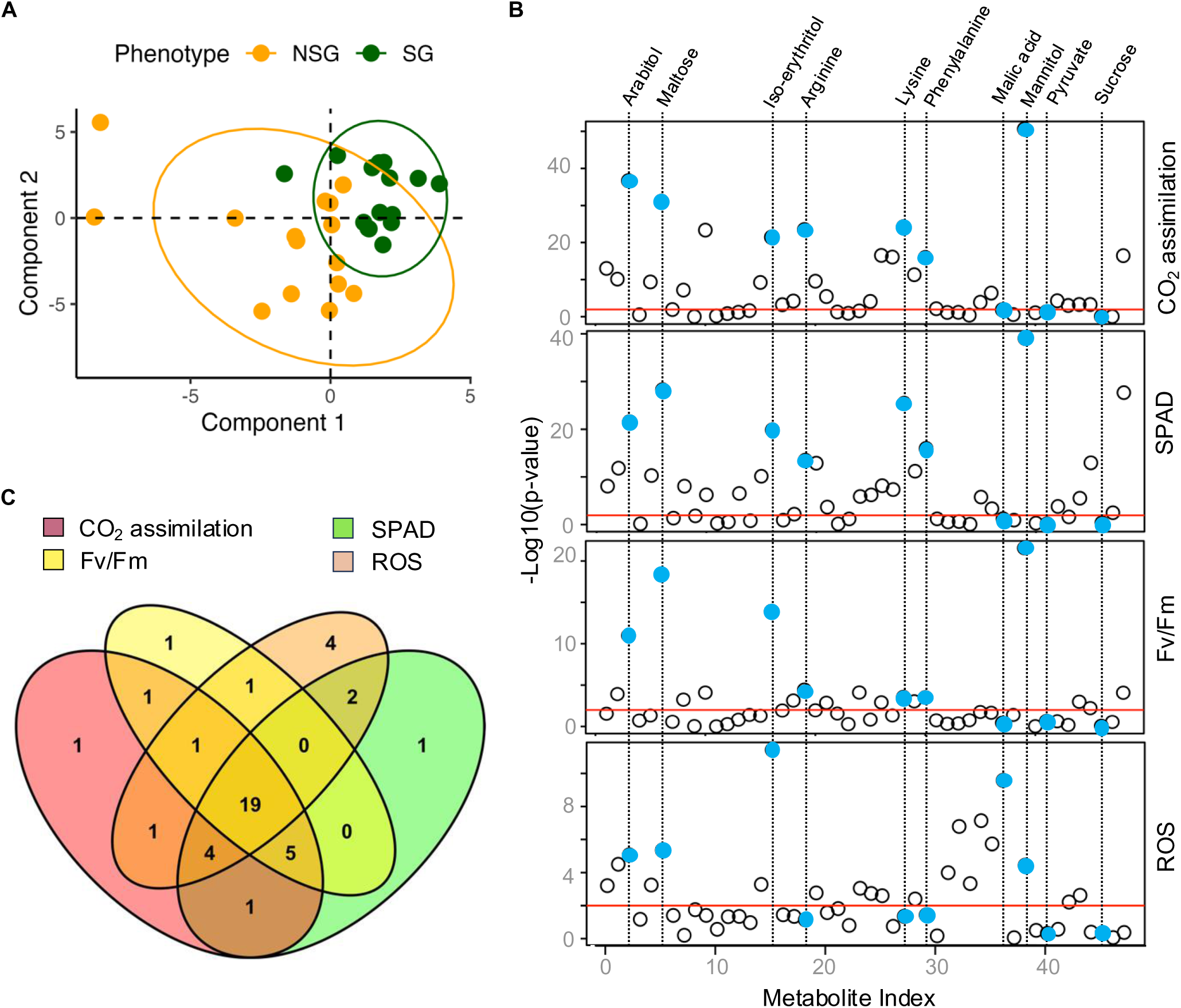
Characterization of primary metabolome associated with senescence. **A**, Partial least square discriminant analysis of primary metabolome of staygreen (SG, green circles) and non-staygreen (NSG, orange circles) inbred lines. Orange and green ovals represent the confidence interval (95%) for the staygreen and non-staygreen inbred lines, respectively. **B**, Marginal analysis of primary metabolites (*x*-axis) with CO_2_ assimilation, chlorophyll content measured by SPAD, Fv/Fm, and reactive oxygen species (ROS). Light blue ovals represent specific metabolites indicated at the top. The red line represents the level of significance (p ≤ 0.01). **C**, Venn diagram shows the number of metabolites associated with different physiological traits.

To examine the association between temporal changes in metabolites and senescence, we performed a marginal analysis that used developmental change in a physiological trait and metabolite abundance as outcome and predictor variables, respectively. Of the 48 detected primary metabolites, 42 (87.5%) showed significant association with at least one physiological trait, and 19 (39.6%) metabolites were significantly associated with all four traits (Figure 3, B-C). These metabolites were represented by four groups that included organic acids (n=6, 14%), sugars and sugar derivatives (n=9, 21%), sugar alcohols (n=7, 17%), and amino acids (n=20, 48%) (Supplementary Table S1). This analysis identified the highest number of primary metabolites significantly associated with CO_2_ assimilation (n=33, 68.7%), followed by SPAD (n=32, 66.7%), ROS content (n=32, 66.7%), and Fv/Fm (n=28, 58.3%) (Supplementary Table S1). These data show that, while all four physiological traits capture some of the fundamental changes in metabolic flux represented by the 19 metabolites, unique metabolites associated with individual phenotypes highlight specific physiological processes captured by these traits.

Since the marginal analysis considered each inbred line separately, some of the physiological and metabolic variation captured by this analysis can also be attributed to genotypic differences among inbred lines irrespective of their staygreen phenotype. Therefore, as a complementary approach to identify metabolites associated with the staygreen phenotype, we performed differential accumulation (DA) analysis for all 48 primary metabolites between the groups of staygreen and non-staygreen inbred lines. This analysis identified 14 metabolites, of which 12 were also revealed by the marginal analysis (Supplementary Table S1). It should be noted that DA analysis relies on the overall staygreen phenotype of inbred lines and, therefore, may not account for diverse mechanisms employed by different genotypes to regulate senescence. Together, the results of these two approaches highlight the importance of using both marginal analysis and DA analyses to identify the metabolite associated with a phenotype.

Consistent with the proposed role of sugar alcohols in the alleviation of stress (Williamson et al., 2002), mannitol, arabitol, iso-erythritol, and galactinol were more abundant in non-staygreen inbred lines (Figure 4). Mannitol was also the top metabolite associated with all three traits related to photosynthesis (i.e., CO_2_ assimilation, Fv/Fm, and SPAD) in marginal analysis while iso-erythritol was the top metabolite associated with ROS (Figure 3B). Amino acids including phenylalanine, arginine, leucine, isoleucine, and lysine were more abundant in the leaves of non-staygreen lines, particularly during the later stages, indicating active proteolysis and nitrogen remobilization during senescence (33 DAA) (Figure 4). Among the sugars, sucrose was more abundant in leaves of non-staygreen inbred lines, while lactose and ribose showed higher accumulation in staygreen inbred lines. Finally, α-ketoglutarate and pyruvate, metabolites related to the TCA cycle, were more abundant in staygreen inbred lines, particularly at the later stages of development. Malic acid showed significant association only with ROS consistent with its protective role under stress conditions (Figure 3B).

**Figure 4:**
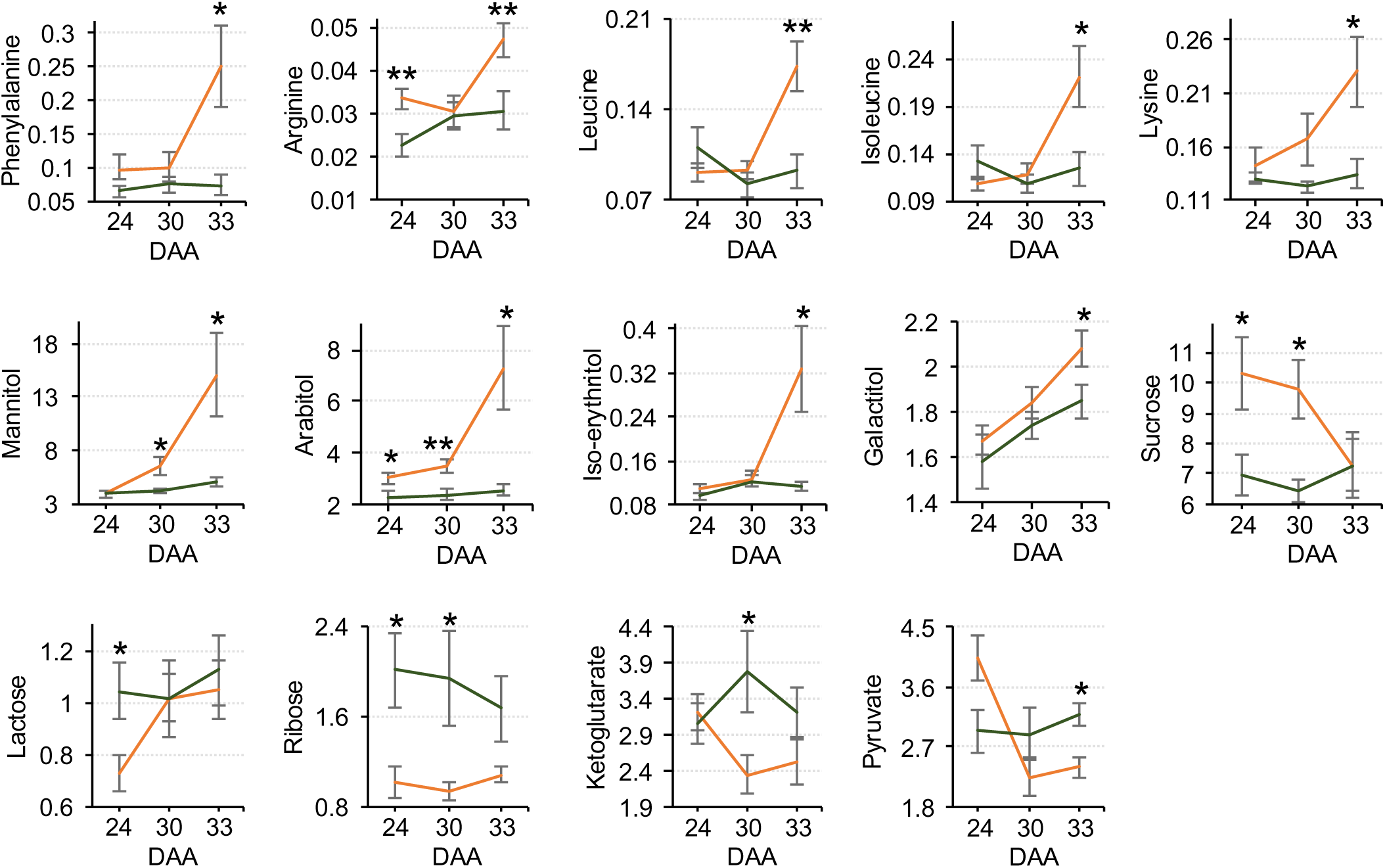
Developmental changes in selected primary metabolites during senescence. Shown here is the average amount of each of the metabolites in the non-staygreen (orange line) and staygreen (green line) inbred lines at three leaf development stages. DAA denotes days after anthesis. One and two asterisks represent the student t-test level of significance p ≤ 0.05 and p ≤ 0.01, respectively.

### Variation in the secondary metabolome

Global analysis of specialized metabolites detected 1,252 mass features. In plant metabolomic experiments, only a fraction of the identified mass features can be annotated because of a lack of reference spectra (da Silva et al., 2015; Alseekh et al., 2021). Therefore, we first used a machine learning approach implemented in CANOPUS (Dührkop et al., 2021) to broadly classify the mass features into distinct chemical classes. This analysis annotated 256 (20.4%) mass features into ten superclasses of organic compounds, of which two most abundant superclasses were represented by organic acids and their derivatives (79, 30.8%) and phenylpropanoids and polyketides (53, 20.4%) (Supplementary Figure S2). Remarkably, PLS-DA based on 1,252 secondary metabolome mass features provided a clear distinction among inbred lines based on the staygreen trait (Figure 5A). These results suggest that the secondary metabolome has a more prominent role in determining the timing and progression of senescence compared to the primary metabolome.

**Figure 5:**
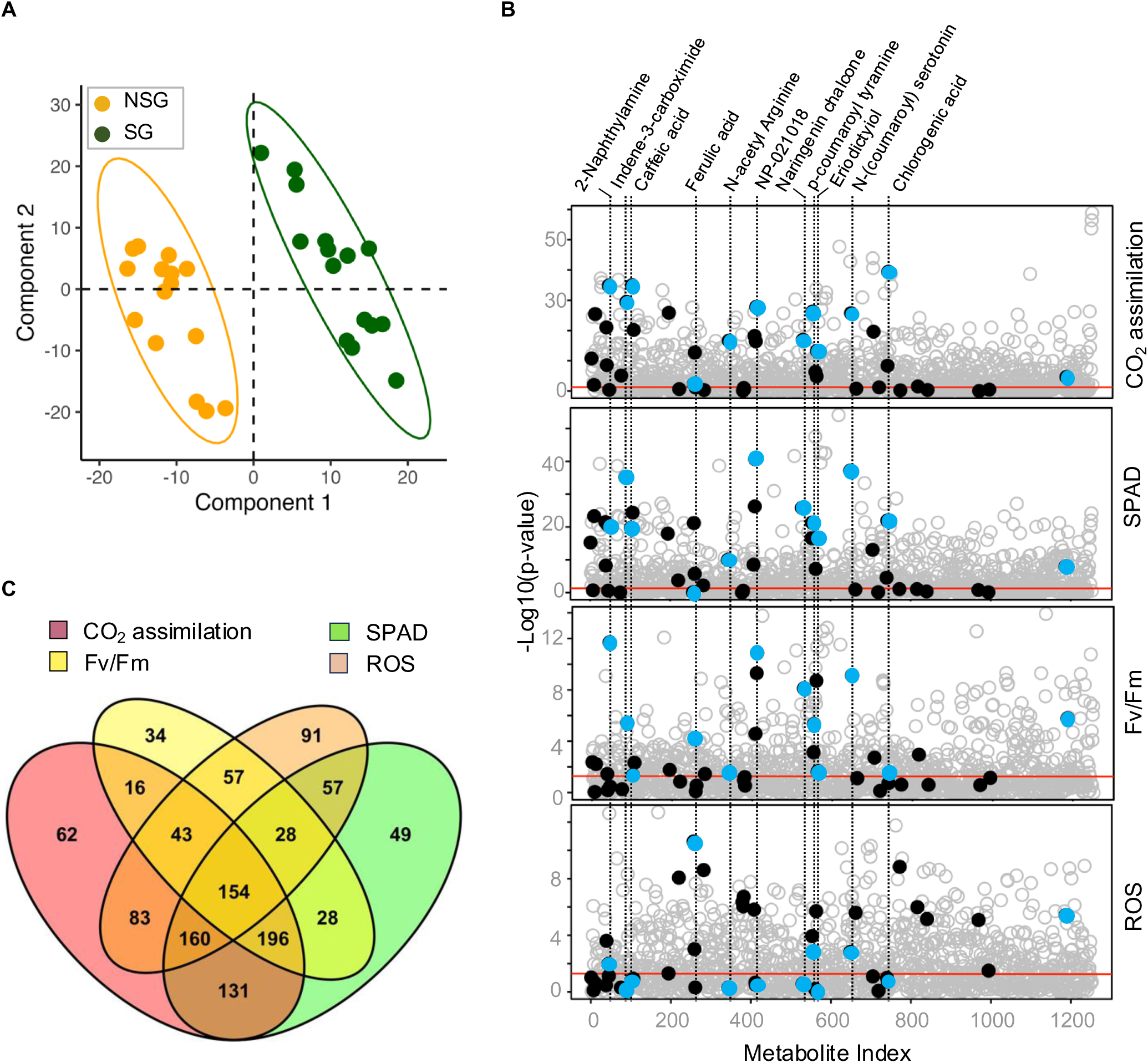
Characterization of secondary metabolome associated with senescence. **A**, Partial least square discriminant analysis of secondary metabolome in staygreen (SG, green circles) and non-staygreen (NSG, orange circles) inbred lines. Orange and green ovals represent the confidence interval (95%) for the staygreen and non-staygreen inbred lines, respectively. **B**, Marginal analysis of secondary mass features (*x*-axis) with CO_2_ assimilation, SPAD, Fv/Fm, and ROS content. Filled circles represent annotated secondary metabolites. Light blue circles represent specific metabolites mentioned at the top. The red line represents the level of significance (p ≤ 0.05). **C**, Venn diagram shows the number of metabolites associated with different physiological traits.

Marginal analysis of the secondary metabolome identified the highest number of mass features significantly associated with CO_2_ assimilation (n=845, 67.5%), followed by SPAD (n=803, 64.1%), ROS content (n=673, 53.7%), and Fv/Fm (n=556, 44.4%) (Figure 5, B-C, Supplementary Dataset S2). Of the 1,252 total mass features, 1,050 (83.8%) showed significant association with at least one physiological trait, and 154 (12.3%) mass features were associated with all four traits (Figure 5C). As a complementary approach to characterize the secondary metabolome associated with senescence, we also performed DA analysis on 1,252 mass features for differential accumulation between staygreen and non-staygreen groups of inbred lines. This analysis identified 201 (16%) mass features of which 189 were also identified by marginal analysis.

Since the identities of the majority of secondary metabolites remain unknown due to the lack of reference spectra (Alseekh et al., 2021), we manually annotated the mass features to identify the corresponding metabolites and understand their biological relevance. For manual curation, we selected the top 100 mass features that showed significant association with each of the four physiological traits and all mass features identified by DA analysis. Of these 601 mass features, we confirmed the identities of 42 (3.3%) secondary metabolites, of which the majority (21, 50%) were phenylpropanoids while the remaining 21 belonged to other groups, including benzaldehydes, terpenes, and amines (Figure 5B, Supplementary Table S2). All 42 annotated secondary metabolites were significantly associated with at least one physiological trait, and 20 of these metabolites differentially accumulated between the staygreen and non-staygreen inbred lines (Supplementary Table S2, Supplementary Figure S3).

Among the annotated secondary metabolites, caffeic acid was associated with all four physiological traits, while chlorogenic acid and ferulic acid, two caffeic acid derivatives, showed significant association with SPAD and CO_2_ assimilation (Figure 5B). However, caffeic acid and chlorogenic acid, not ferulic acid, showed significantly higher accumulation in the leaves of staygreen inbred lines during senescence (Figure 6). Interestingly, ferulic acid was the top metabolite associated with ROS indicating a fundamental role of this metabolite in mitigation of oxidative stress. Among flavonoids, naringenin chalcone and eriodyctiol were significantly associated with CO_2_ assimilation rate, SPAD, and Fv/Fm (Figure 5B), albeit only eriodyctiol showed differential abundance between non-staygreen and staygreen groups of inbred lines during senescence (Figure 6). Among amines, secondary metabolites with diverse functions including osmotic stress response and delaying senescence, 2-Napthylamine, N-(p-coumaroyl) serotonin, and p-coumaroyl tyramine were significantly associated with all four physiological traits (Figure 5B). Both 2-Napthylamine and p-coumaroyl tyramine also showed significantly higher accumulation in the leaves of non-staygreen inbred lines during senescence (Figure 6). Finally, NP-021018 was associated with all four physiological traits and had significantly higher accumulation in staygreen inbred lines.

**Figure 6:**
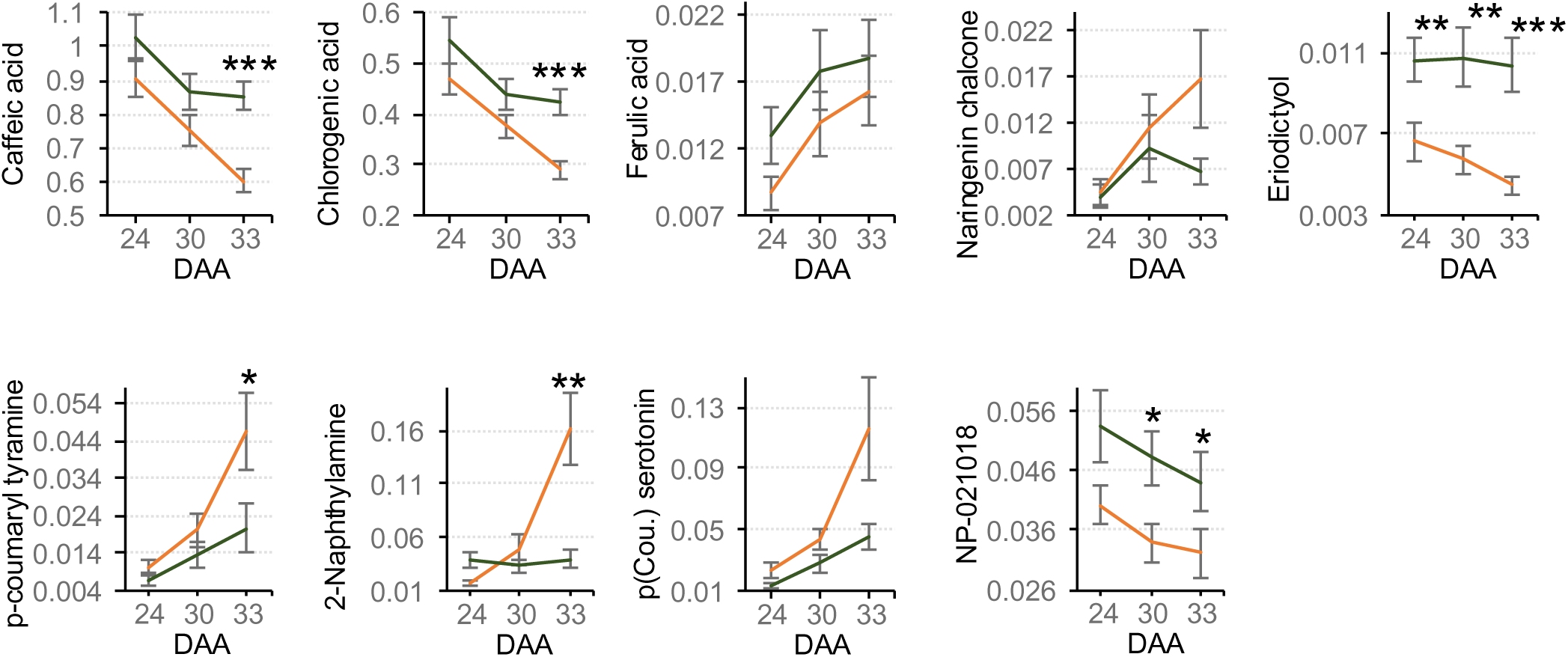
Developmental changes in selected secondary metabolites during senescence. Shown here is the average amount of each of the metabolites in staygreen (green lines) and non-staygreen (orange lines) inbred lines at three leaf development stages. One, two, and three asterisks represents the student t-test level of significance at p ≤ 0.05, p ≤ 0.01, and p ≤ 0.001, respectively.

### Enrichment analysis identifies specific pathways underlying senescence

To identify the metabolic pathways that are differentially employed by staygreen and non-staygreen inbred lines, we performed pathway enrichment analysis using all 1,252 mass features. This analysis identified four metabolic pathways including phenylpropanoid biosynthesis, flavonoid biosynthesis, biosynthesis of various plant secondary metabolites, and cyano-amino acid metabolism that differentiated staygreen and non-staygreen inbred lines (Figure 7, Supplementary Figure S4). Identification of phenylpropanoid and flavonoid biosynthesis pathways complements the discovery of flavonoids and caffeic acid derivatives in statistical analyses of secondary metabolites (Figure 5B, 6). Detection of the cyano-amino acid metabolism pathway underscores the role of cyanide-conjugated amino acid metabolites in determining the staygreen trait. Collectively, these findings highlight key metabolic pathways that play a crucial role in determining the staygreen trait.

**Figure 7:**
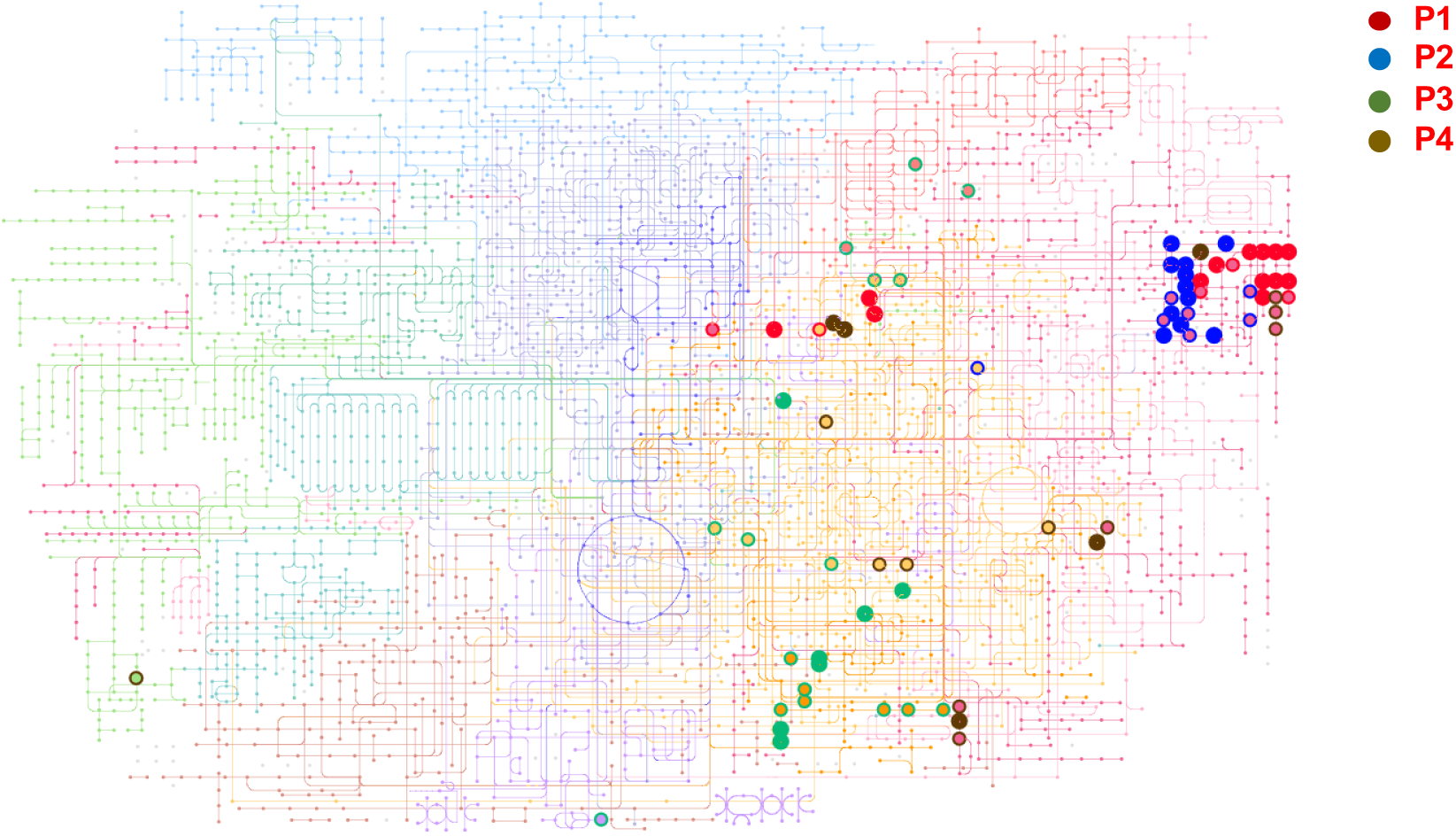
Pathway enrichment analysis of secondary mass features. P1, P2, P3, and P4 represent the phenylpropanoid biosynthesis pathway, flavonoid biosynthesis pathway, biosynthesis of various plant secondary metabolites pathway, and cyano-amino acid metabolism pathway, respectively.

### Identification of genes and networks regulating the senescence-associated metabolome

Metabolites are internal phenotypes that represent the functional readout of cellular biochemistry and, therefore, offer a reliable link between the genome and the external plant phenotypes. Structure-function investigations of enzymes and metabolic pathways are commonly used to derive functional annotations for these biochemical processes and their associated enzymes (Monaco et al., 2013). To identify genes responsible for variation in the metabolome, we set out to identify enzymes catalyzing biosynthesis of each of the 84 primary and secondary metabolites that showed significant association with at least one physiological phenotype and the genes encoding for these enzymes. This approach identified 186 genes encoding for distinct enzymes catalyzing the biosynthesis of 33 (39%) metabolites including 24 primary and nine secondary metabolites (Supplementary Dataset S3). Through extensive literature searches, we shortlisted 56 novel candidate genes expressed in mature leaves that encode for enzymes responsible for the biosynthesis of these metabolites (Table 2)(Thomazeau et al., 2001; Diebold et al., 2002; Joshi et al., 2006; Martin et al., 2006; Todd et al., 2008; Holding et al., 2010; Liu et al., 2019).

**Table 2:** Putative genes encoding enzymes upstream of metabolites.

| Metabolites | Enzymes | Putative genes |
| --- | --- | --- |
| Mannitol | Mannitol phosphatase | Zm00001eb113090 |
| Lysine | Diaminopimelate decarboxylase | Zm00001eb406820 |
| Arginine | Argininosuccinate lyase | Zm00001eb396370 |
| Leucine | Leucine transaminase | Zm00001eb404340, Zm00001eb074240, Zm00001eb022930, Zm00001eb017520, Zm00001eb009540 |
| Phenylalanine | Arogenate dehydratase | Zm00001eb227480, Zm00001eb383660 |
| Methionine | Methionine synthase | Zm00001eb032290 |
| Ketoglutarate | Isocitrate dehydrogenase | Zm00001eb434270, Zm00001eb296460 |
| Asparagine | L-aspartate:L-glutamine amido-ligase | Zm00001eb013430, Zm00001eb380080, Zm00001eb396990 |
| Citric acid | Citrate synthase | Zm00001eb206090, Zm00001eb206100 |
| Aspartic acid | Cyanoalanine nitrilase | Zm00001eb185550 |
| Alanine | Glutamate--pyruvate aminotransferase | Zm00001eb028030 |
| Histidine | Histidine dehydrogenase | Zm00001eb059830 |
| Tyrosine | Arogenate dehydrogenase | Zm00001eb383660 |
| Succinic acid | Succinyl-CoA synthetase | Zm00001eb107590, Zm00001eb185030, Zm00001eb325420 |
| Shikimic acid | Shikimate dehydrogenase | Zm00001eb131900, Zm00001eb410660 |
| Glycine | Glycine hydroxymethyltransferase | Zm00001eb146170 |
|  | Glycine transaminase | Zm00001eb034260, Zm00001eb118240, Zm00001eb298440 |
|  | Threonine aldolase | Zm00001eb077640 |
| Glutamine | Glutamine synthetase | Zm00001eb009090, Zm00001eb190340, Zm00001eb253820, Zm00001eb432580 |
| Cysteine | Cysteine synthase | Zm00001eb032380, Zm00001eb127210 |
| Glucose-6-Phosphate | Glucose-6-phosphate isomerase | Zm00001eb059230, Zm00001eb101090, Zm00001eb315420 |
| Tryptophan | Tryptophan synthase | Zm00001eb060880, Zm00001eb301540 |
| Fumaric acid | Succinate dehydrogenase | Zm00001eb173990 |
| Proline | Pyrroline-5-carboxylate reductase | Zm00001eb141220 |
| Threonine | Threonine synthase | Zm00001eb294790 |
| Serine | 3-phosphoserine aminotransferase | Zm00001eb004390 |
| Adenosine 5'-monophosphate | 4-coumarate--CoA ligase | Zm00001eb233720 |
| Chlorogenic acid | NADPH:oxygen oxidoreductase | Zm00001eb150550, Zm00001eb291460 |
| Tyramine | Tyrosine decarboxylase | Zm00001eb292480 |
| Ferulic acid | Caffeate O-methyltransferase | Zm00001eb172420 |
| Eriodictyol | Flavonoid 3' hydroxylase | Zm00001eb245960 |
| Uracil | Uridine nucleosidase | Zm00001eb039560 |
| Kaempferol | Flavonol synthase | Zm00001eb256020 |
| Cynaroside | Uridine diphosphate glycosyltransferase | Zm00001eb380880 |
| Naringenin chalcone | Chalcone synthase | Zm00001eb198030 |

To generate additional evidence for the role of these genes in senescence, we used published gene expression datasets obtained from three distinct genotypes (Sekhon et al., 2019; Kumar et al., 2023) to construct a co-expression network of genes associated with leaf senescence. This network identified a large sub-network of 4193 genes and two smaller sub-networks of 44 and 37 genes (Figure 8A). Remarkably, of the 56 genes encoding for senescence-associated metabolites, 10 were present in the larger network, and one gene was represented in a smaller network (Supplementary Dataset S3). Additionally, six genes reported in previous reverse genetics studies (Fahnenstich et al., 2007; Slewinski et al., 2009; Urano et al., 2014; Li et al., 2015; Yu et al., 2019; Zhang et al., 2019) and 13 candidate genes identified from genome-wide association (GWA) analysis of natural variation for senescence (Sekhon et al., 2019; Kumar et al., 2023) were also represented in these networks. In summary, linking metabolome to genome provided a core set of novel genes, and network analysis provided additional proof of evidence for the role of a subset of these genes in senescence.

**Figure 8:**
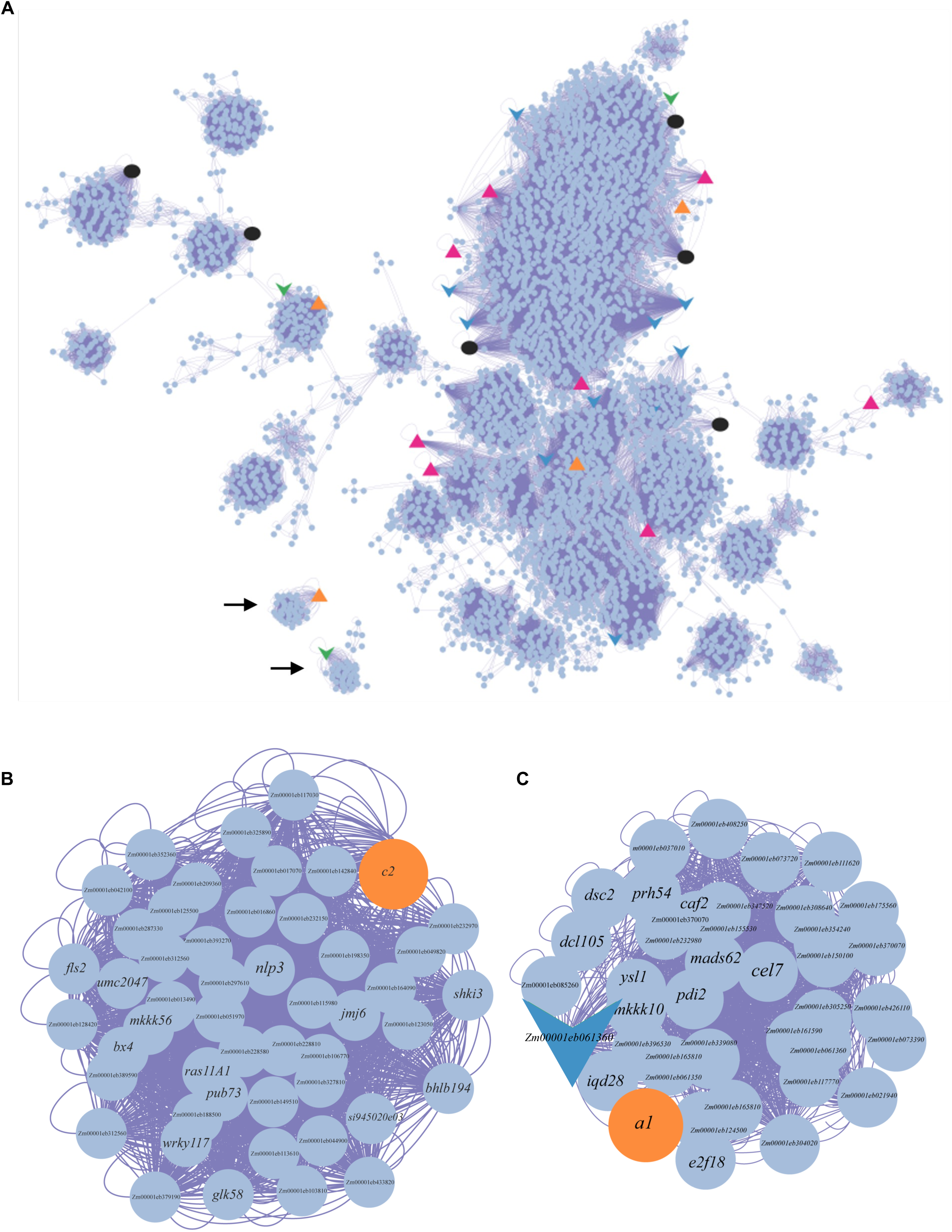
Meta-analysis of candidate genes associated with senescence in maize. **A**, One large (top) and two small (marked by arrows) co-expression networks associated with senescence with genes identified from other omics analysis indicated by different shapes. Pink and orange triangles represent candidate genes responsible for the biosynthesis of primary and secondary metabolites identified in the current study, respectively, black circles represent previously known genes regulating senescence in maize, and blue and green arrowheads denote candidate genes identified by analysis of natural variation for natural and source-sink regulated senescence, respectively. **B**, A module comprised of *colorless2* (*c2*) (orange circle) and other interacting partners. **C**, A module comprised of *anthocyaninless1* (*a1*) (orange circle), a candidate gene identified by analysis of natural variation for senescence (blue arrowhead), and other interacting partners.

### Genetic analyses support the role of naringenin chalcone and eriodictyol in senescence

Identification of naringenin chalcone and eriodictyol as important secondary metabolites prompted us to ask if the genes involved in regulating these flavonoids have a role in senescence. The flavonoid pathway is well characterized and structural genes responsible for the synthesis of key enzymes have been identified in maize and Arabidopsis (Dooner et al., 1991; Saito et al., 2013). In this pathway, chalcone synthase encoded by *colorless2* (*c2*) catalyzes the synthesis of naringenin chalcone from 4-Coumaroyl-CoA while dihydroflavonol 4-reductase encoded by *anthocyaninless1* (*a1*) catalyzes the conversion of eriodictyol to luteoferol (Figure 9A) (Schröder, 1997; Lo and Nicholson, 1998; Sharma et al., 2011; Yang et al., 2017). Remarkably, both *c2* and *a1* genes were mapped on the senescence co-expression network (Figure 8), further supporting the role of these metabolites in regulating the staygreen trait. The module containing *c2* also includes other interesting candidate genes including *wrky117* transcription factor shown to promote senescence in Arabidopsis, *flavanol synthase2* (*fls2*) involved in abiotic stress tolerance (Figure 8B) (Falcone-Ferreyra et al., 2012; Yu et al., 2021). The module containing *a1* also included several novel genes including *mads62* and *e2f18* transcription factors, and a novel candidate gene identified in a previous analysis of natural variation for source-sink regulated senescence (Figure 8C). These modules present novel information relevant to the regulation of flavonoids in maize leaves and their role in leaf senescence.

**Figure 9:**
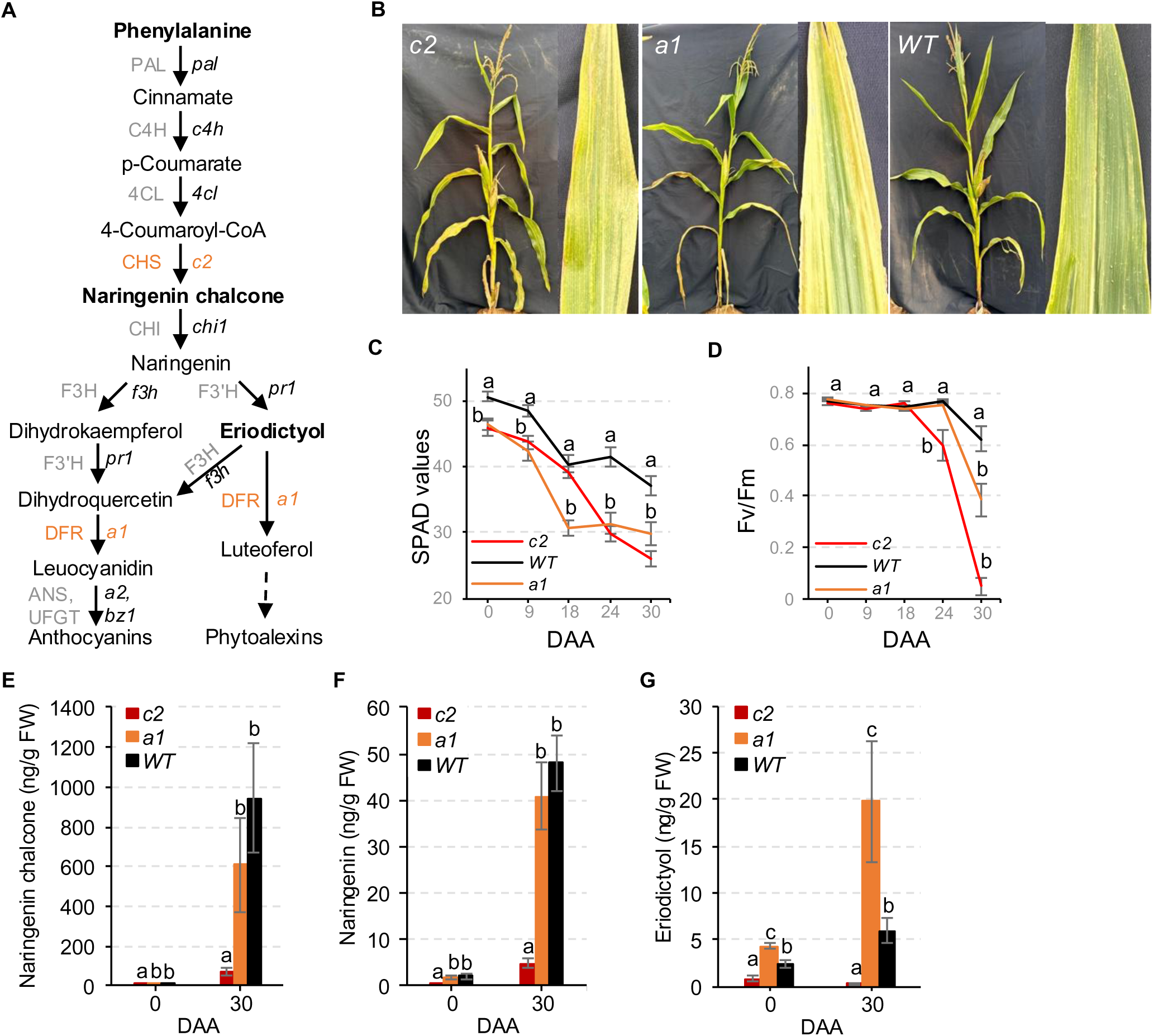
Genetic analysis of the role of naringenin chalcone and eriodictyol in maize leaf senescence. **A**, Schematic showing the flavonoid biosynthetic pathway. The enzymes (PAL, phenylalanine ammonia lyase; C4H, cinnamate 4-hydroxylase; 4CL, 4-coumarate CoA ligase; CHS, chalcone synthase; CHI, chalcone isomerase; F3H, flavanone 3-hydroxylase; F3’H, flavonoid 3’-hydroxylase; DFR, dihydroflavonol reductase; UFGT, UDP-glucose flavonoid 3-O-glucosyl transferase; ANR, anthocyanidin reductase) are mentioned on left and the genes encoding these enzymes are provide on right in italicized font. Metabolites denoted by boldfaced font were identified in our metabolome analysis. Genes of interest are highlighted in orange text. **B**, Visual manifestation of leaf senescence in mutant and wild-type plants at 24 DAA. **C-D**, SPAD and Fv/Fm values, respectively, of the top ear-bearing leaf in mutant and wild-type plants. Alphabets represent the level of significance based on Tukey HSD test at p ≤ 0.05. **E-G**, Concentration of naringenin chalcone, naringenin, and eriodictyol, respectively, in mutant and wild-type plants at 0 and 30 DAA. Alphabets represent the level of significance based on Tukey HSD test at p ≤ 0.05.

To further examine the role of naringenin chalcone and eriodictyol in senescence, we evaluated maize *c2* and *a1* transposon insertional mutant plants. Visually, both *c2* and *a1* mutant plants showed early senescence after flowering compared to their wild-type counterparts as evident from the yellowing of leaves (Figure 9B). Both *c2* and *a1* mutant plants had lower chlorophyll content throughout the post-flowering period as indicated by SPAD data and significantly decreased Fv/Fm values during later stages of the post-flowering period (Figure 9, C-D). Consistent with the role of chalcone synthase in the metabolic pathway, the *c2* mutants accumulated a lower amount of naringenin chalcone, naringenin, and eriodictyol (Figure 9, E-G). As expected, due to the loss of dihydroflavonol 4-reductase activity, *a1* mutant plants had significantly high eriodictyol content compared to wild-type plants suggesting that conversion of eriodictyol to downstream compounds is vital for timely senescence (Figure 9, E-G). Collectively, these genetic data support the role of the identified flavonoids in maize leaf senescence.

The flavonoid biosynthetic pathway and the associated enzymes are highly conserved in maize and Arabidopsis (Falcone Ferreyra et al., 2012) (Figure 9A), therefore, we asked if the identified flavonoids have a conserved role in senescence across monocots and dicots. The phenotypic evaluation showed early visual senescence and lower Fv/Fm values in both *chs* and *dfr* mutants supporting the role of these respective enzymes in senescence (Figure 10, A-B). Interestingly, both mutants also accumulated higher ROS in leaves indicating a combinatorial role of flavonoid accumulation and quenching of ROS by these flavonoids in regulation of leaf senescence (Figure 10C). Consistent with the role of chalcone synthase in the metabolic pathway, the *chs* mutants accumulated a lower amount of naringenin chalcone and naringenin and no detectable levels of eriodictyol (Figure 10D). Interestingly, the *dfr* mutants also had a lower accumulation of naringenin chalcone and naringenin but did not show any change in eriodictyol (Figure 10D). This observation can be explained by the diversion of overall flux from the pathway upstream of naringenin chalcone to other metabolites due to a lack of functional downstream enzymes. In summary, these data indicate that the flavonoids identified in the study have a conserved role in leaf senescence in these two distinct species.

**Figure 10:**
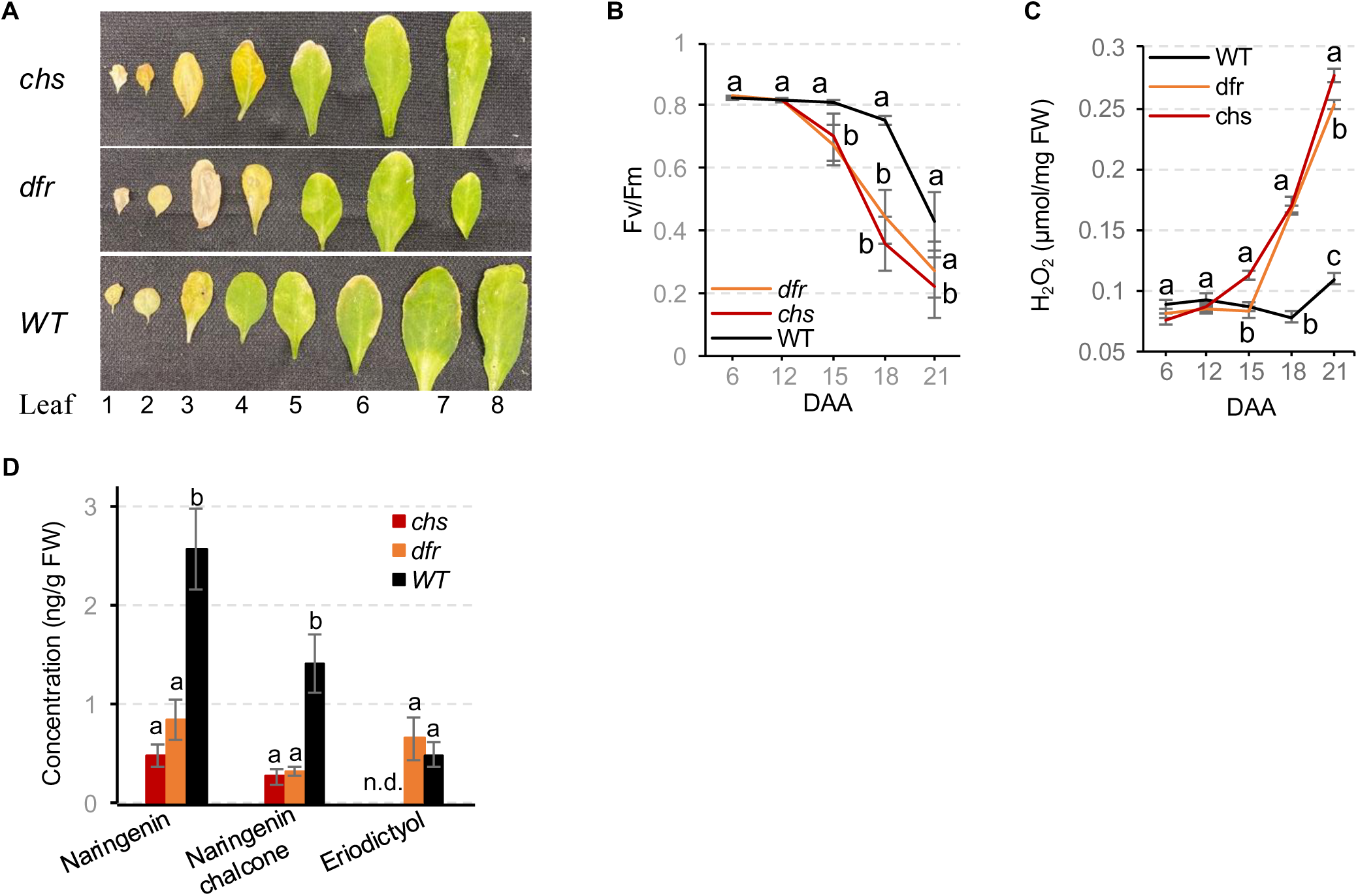
Genetic analysis of the role of naringenin chalcone and eriodictyol in Arabidopsis leaf senescence. **A**, Visual manifestation of leaf senescence in mutant and wild-type plants at 15 DAA. **B**, Fv/Fm values of the fourth leaf in mutant and wild-type plants. Alphabets represents the level of significance based on Tukey’s HSD test at p ≤ 0.05. **C**, ROS content in the fourth leaf in mutant and wild-type plants. Alphabets represents the level of significance based on Tukey’s HSD test at p ≤ 0.05. **D**, Concentration of naringenin chalcone, naringenin, and eriodictyol in chalcone synthase (*chs*) and dihydroflavonol reductase (*dfr*) mutant plants and wild-type plants at 15 DAA (nd, not detected). Alphabets represents the level of significance based on Tukey’s HSD test at p ≤ 0.05.

## DISCUSSION

Leaf senescence is accompanied by large-scale changes in physiological and metabolic phenotypes, although the nature of the changes and the interplay between these phenotypes remains largely unknown. Numerous studies have examined a few specific physiological and metabolic phenotypes to generate valuable insights into the biological underpinnings of senescence (Wingler and Roitsch, 2008; Watanabe et al., 2013; Liebsch et al., 2022; Zhu et al., 2022; Perez-Fons et al., 2023). However, senescence is a dynamic process involving global shifts in diverse phenotypes and only partial information can be generated by studying a snapshot of one phenotype at a time. In this study, we elucidated the dynamics of leaf metabolome and distinct physiological phenotypes during the onset and progression of senescence in a set of diverse maize inbred lines. Combinatorial analysis of temporal changes in physiological phenotypes and metabolome identified novel primary and secondary metabolites associated with leaf senescence. We annotated the key metabolites and assigned them to respective pathways to highlight the role of specific metabolic networks and identify the underlying enzymes and genes. Reverse genetic analysis of genes regulating two distinct metabolites in flavonoid biosynthesis pathways in maize and Arabidopsis confirmed the role of identified metabolites in senescence.

Senescence is a complex process and the lack of consensus on the effectiveness of strategies for quantitative measurement of the phenotypic manifestations of this process poses a major obstacle to understanding the genetic underpinnings. For instance, plants manifest staygreen in multiple ways ranging from cosmetic staygreen wherein the delayed senescence does not translate to increased photosynthetic assimilation to functional staygreen resulting in higher carbon yield (Thomas and Howarth, 2000). Therefore, the choice of phenotyping method is crucial for discerning the nature of staygreen phenotype and uncovering the underlying specific genetic networks. A comparison of five different physiological traits showed that CO_2_ assimilation, SPAD, and Fv/Fm produced a similar grouping of inbred lines consistent with empirical data on the staygreen trait for many of these inbred lines. Given that CO_2_ assimilation directly measures photosynthetic assimilation, similar clustering of staygreen inbred lines indicates the functional nature of the staygreen trait observed in these lines. However, ROS content and C/N ratio failed to provide distinct clustering, indicating that variation for these more intrinsic processes was conserved during the selection of the staygreen phenotype. Notably, the lack of a clear association of the C/N ratio with staygreen phenotype in these inbred lines shows extant variation for C partitioning and N remobilization in maize germplasm. Finally, the inability of a single timepoint of phenotypic measurement to correctly classify the inbred lines for the staygreen phenotype highlights the importance of temporal measurement for phenotypic and genetic analysis of senescence.

The emergence of sugars, sugar alcohols, and amino acids as major groups of primary metabolites highlights the significance of C and N metabolism and altered source-sink relationship during senescence. Senescing leaf undergoes a switch from a C capture and transport phase dominated by anabolic processes to an N remobilization phase dominated by catabolic activities (Schippers et al., 2015; Havé et al., 2016). Deceased sink demand after grain fill results in disruption of sugar partitioning and accumulation of soluble sugars in the leaves that play a key role in senescence signaling (Sekhon et al., 2012; Kumar et al., 2019; Kumar et al., 2023). Higher accumulation of sugar alcohols in non-staygreen leaf cells during senescence indicates an important albeit poorly understood role of these sugar derivatives in leaf senescence. There is, however, an indirect evidence for such a role as mannitol treatment has been shown to suppress the spike in sucrose and glucose content and inhibit flower senescence (Ichimura et al., 2016). Likewise, erythritol has been reported to delay senescence in mice and human cells likely by promoting hexose breakdown and pyruvate production (Yokoi et al., 2023). Given the role of hexoses in senescence signaling through HXK1-mediated suppression of cytokinin (Kumar et al., 2023), these sugar alcohols may suppress such signaling and slow senescence.

Increased content of several amino acids in non-staygreen inbred lines indicates enhanced N remobilization, and this observation is consistent with the low N content in leaves of non-staygreen inbred lines (Supplementary Figure S5). Our previous studies have highlighted the role of proteolysis mediated by two distinct cysteine proteases in determining the senescence phenotype (Sekhon et al., 2019; Kumar et al., 2023). Leaf N remobilization during senescence contributes 50% to 90% of grain N content in cereals thus underscoring the importance of the nitrogen remobilization efficiency to grain quality (Kichey et al., 2007; Masclaux-Daubresse et al., 2008). Conversely, the application of N fertilizers and release of excess N to the environment accounts for an estimated 6% of the entire US greenhouse gas emissions and 55% of agricultural emissions (EPA, 2024). Therefore, the identification of molecular components of N remobilization during senescence holds the key to improving grain quality and N economy. A higher concentration of metabolites related to the TCA cycle including pyruvate and α-ketoglutarate indicates active cellular metabolism and energy generation in staygreen inbred lines. While such cellular bioenergetics could be seen as an outcome of the active cellular function, the regulatory role of these metabolites from the TCA cycle in senescence needs to be explored. Interestingly, α-ketoglutarate has been shown to delay heat-induced leaf senescence by mitigating oxidative stress in perennial ryegrass and promoting photosynthesis and plant growth in poplar (Lei and Huang, 2022; Liu et al., 2024).

The emergence of secondary metabolisms as the major hallmark of the senescence metabolome underscores the importance of these compounds in alleviating stress and maintaining cellular function during senescence. A wide variety of secondary metabolites are derived from phenylalanine, tryptophan, and tyrosine and, while all three groups are represented in the senescence-associated secondary metabolome, phenylpropanoids emerged as the predominant group. An increase in phenylpropanoids, mainly organic acids and flavonoids, coincides with high ROS content suggesting a role of these metabolites in mitigating oxidative stress during senescence. Consistent with these findings, the exogenous application of caffeic acid, ferulic acid, and certain flavonoids has been shown to delay senescence by mitigating oxidative stress (Sgarbossa et al., 2015; Fan et al., 2022; Huang et al., 2022). While the flavonoids have been previously linked with senescence (Li et al., 2017; Xue et al., 2021), functional characterization of genes involved in the regulation of naringenin chalcone and eriodictyol in maize and Arabidopsis indicates a conserved role of these metabolites in leaf senescence in monocots and dicots. While the exact mechanism(s) through which these secondary metabolites impact senescence needs to be investigated, higher ROS content in non-staygreen inbred lines and Arabidopsis mutants lacking these metabolites support their role as ROS quenchers. Alternatively, sugar-flavonoid conjugates may compromise the signaling properties of hexoses and, therefore, delay senescence.

While metabolites arguably provide the most reliable link between the genome and external phenotypes, linking metabolites to the genes essential for the regulation and manifestation of senescence has lagged. Linking senescence-associated metabolome to the genome in the current study identified 56 genes catalyzing the synthesis of 33 metabolites that are reasonable targets to study senescence. Remarkably, meta-analyses of the genes linked with senescence metabolome in the current study supported the role of 11 of these 56 candidate genes in senescence. Validation of these novel genes in senescence needs further investigation using reverse genetics and functional analyses. The lack of annotation of over 93% of secondary metabolites is a substantial knowledge gap that hinders the identification of underlying processes and genes and offers exciting opportunities for future studies. Besides improving analytical approaches, this goal can be accomplished through genomic studies including metabolite quantitative trait locus and metabolite-wide association analyses to discover the genes regulating the flux of these metabolites.

In conclusion, the present study highlights the need for a paradigm shift in phenotyping approaches to account for the dynamic nature of senescence to characterize the genetic architecture of this complex process. The identification of 84 primary and secondary metabolites provides a set of valuable targets for future studies to discover the genes and mechanisms underlying senescence phenotype. Identification of 56 novel candidate genes regulating the biosynthesis of these metabolites, a fifth of which are also supported by a meta-analysis of additional genomic studies, offers opportunities to enhance the genetic architecture of senescence. Functional validation of eriodictyol and naringenin indicates a novel role of the secondary metabolites in senescence and further demonstrates the value of metabolic analyses in identifying the underlying genes.

## MATERIAL AND METHODS

### Plant materials

Maize inbred lines (Table 1) were grown at the Clemson University Simpson Research Farm (CUSRF), Pendleton, South Carolina, USA in a randomized complete block design with three replications. Each 11.12 m^2^ plot consisted of three 4.88 m long rows with row and plant spacing of 76 cm and 20 cm apart, respectively. All the plants in a plot were open-pollinated when 80% of plants showed silk emergence and this stage was marked as 0 DAA. Physiological data was recorded on 6-10 plants in each plot. All physiological data were collected on the adaxial side of the leaf blade approximately six inches from the base of the leaf while carefully avoiding the mid-rib. Fv/Fm data were collected on dark-adapted leaves 1 hr after sunset using a Pocket PEA fluorimeter (Hansatech Instruments) by exposing the leaves to a light intensity of 3500 μmol m^−2^ s^−1^ for 1 s. CO_2_ assimilation was measured in the morning (9 am - 12 pm) with LI-6800 portable photosynthesis systems (LI-COR Inc., Lincoln, NE, USA) keeping air humidity at 60%, leaf temperature close to ambient conditions (25°C to 32°C), the light intensity of 2000 μmol m^−2^ s^−1^, and CO_2_ concentration of 500 ppm. SPAD data was collected in the morning (7 am – 9 am) using SPAD 502 chlorophyll meter (Konica Minolta Inc., Osaka, Japan). For each plot and stage, the whole leaves at the primary ear-bearing node excluding ligule, midrib, and sheath were harvested and pooled from two competitive randomly chosen plants between 10 am to 2 pm, immediately frozen in liquid N_2_, and stored in −80°C freezer. For all metabolic analyses, the leaf samples were ground to a fine powder under liquid nitrogen with a mortar and pestle, transferred to 50 ml falcon tubes, and stored in a −80°C freezer for future analyses.

For maize mutant analysis, genetic stocks in W23 background including M142A (*A1*; *A2; b1*; *C1*; *C2*; *pl1*; *R1-r*), M142B (*a1*; *A2; b1*; *C1*; *C2*; *pl1*; *R1-r*), M142H (*A1*; *A2; b1*; *C1*; *c2*; *pl1*; *R1-r*) were kindly provided by the Maize Genetics Cooperation Stock Center (USDA Agricultural Research Service, University of Illinois, Urbana, IL). These stocks were grown at the CUSRF, Pendelton, South Carolina, USA during the summer of 2024 in a randomized complete block design with three replications. Each replication consisted of a 3.71 m^2^ single-row plot of one 4.88 m long row with row and plant spacing of 76 cm and 20 cm apart, respectively. Physiological data, tissue collection, and tissue preparation for metabolic analyses were performed as described for the maize inbred lines experiment. Physiological data was recorded on ten representative plants in each replication.

For Arabidopsis mutant analysis, genetics stocks in L*er* background including CS85 (*chs*), CS84 (*dfr*), and CS20 (*CHS*; *DFR*) were obtained from the Arabidopsis Biological Resource Center (https://www.arabidopsis.org/). Arabidopsis plants were grown in the growth room as described (Sekhon et al., 2019). The fourth true leaf was dark-adapted by placing a clamp for 10 min and Fv/Fm data was recorded with a Pocket PEA fluorimeter (Hansatech Instruments) by exposing the leaves to a light intensity of 3500 μmol m^−2^ s^−1^ for 1 s. Fv/Fm data was recorded on eight plants from each genotype. For metabolic analysis, all rosette leaves from three individual plants from each genotype were harvested, immediately frozen in liquid N_2_, ground to fine powder with a mortar and pestle, and stored in a −80°C freezer for future analyses.

### Carbon and Nitrogen content measurement

Finely ground frozen leaf tissue was freeze dried and 3-5 mg tissue was aliquoted to tin capsules for the analyses. Total carbon and nitrogen were measured using the combustion method (Luo et al., 2017) at the University of New Mexico Center for Stable Isotope facility.

### ROS measurement

H_2_O_2_ concentration was measured using the Amplex Red H_2_O_2_/Peroxide Assay kit (Invitrogen) (Zhang et al., 2021). Briefly, 30 mg of finely ground frozen leaf tissue was suspended into 200 μl H_2_O_2_ extraction buffer, centrifuged at 12,500 rpm for 20 min at 4°C, and the supernatant was transferred to the new tubes for quantification. Fluorescence was measured with a microplate reader using excitation of 530 nm and detection at 590 nm.

### Metabolome analysis

To extract metabolites, 100 mg of frozen ground leaf tissue was weighed into 2 ml tubes and homogenized with 1.0 ml methanol using a bead homogenizer. The homogenized samples were centrifuged at 6,000 rpm for 2 min, and the supernatant was collected and stored at −80C for further analyses. For metabolomic analyses, 0.5 ml of ice-cold chloroform and 0.5 ml of cold water were added to 0.5 ml of the supernatant. The mixture was centrifuged at 6000 rpm for 2 min, and the top methanol-water phase was stored in a −80°C freezer for further analyses.

Untargeted primary metabolomics to capture compounds of central C and N pathways analysis was performed using gas chromatography-mass spectrometry (GC-MS) as described (Sandhu et al., 2023). Briefly, the chloroform-partitioned methanol-water extract was derivatized using myristic acid-d27 and ribitol as internal standards and the derivatized extracts were dried under a nitrogen gas stream and subjected to methoximation and silylation. The metabolites were separated and analyzed using a single quadrupole mass spectrometer and gas chromatography (Agilent 5975). Quality control samples were prepared by mixing equal aliquots from all samples and run after every 15 samples. Mass spectra were prepared for peak selection, alignment, and integration using MS-Dial. The annotation of principal metabolites was based on the Kovats retention index, mass fragmentation matches with a minimum EI similarity score of 80%, and manual curation to confirm the library hits.

The untargeted secondary metabolome analysis was performed using ultrahigh pressure liquid chromatography-tandem mass spectrometry (UHPLC-MS/MS) using an Ultimate 3000 UHPLC connected to an Orbitrap Fusion Tribrid mass spectrometer with a heated electrospray ionization (HESI) source as published (Sandhu et al., 2023). Briefly, a 150 μL aliquot of the chloroform-partitioned methanol-water extract was combined with 150 μL of 13C6-resveratrol (0.5 μg mL^-1^) as internal standard and 4 μL of the mixture was separated on an Acclaim PepMap 100 C18 column (150 × 1 mm, 3 μm) maintained at 32°C. Pooled quality control samples prepared by pooling all samples and method blanks were run after every 15 samples. The mobile phases A and B consisted of 0.05% formic acid in water and acetonitrile, respectively. The solvent gradient started at 5% B, increased to 95% B, and returned to 5% B. The HESI ion source was operated in positive ionization mode and data-dependent MS2 mass spectra were collected. The MS1 scan data was collected in an Orbitrap with an intensity threshold and dynamic exclusion protocol. MS2 fragmentation was performed using stepped higher-energy collisional dissociation values and the mass spectra were collected in the Orbitrap. For compound identification, the pooled sample was analyzed in negative ionization mode using data-dependent MS2 and the UHPLC–MS/MS spectra were processed using Compound Discoverer v3.1. Compounds were annotated based on accurate mass (±5 ppm) and characteristic fragmentation patterns, online libraries (MoNA, mzCloud, Flavanoid Search, HMDB), literature search, and compound class prediction of CANOPUS and CSI: Finger ID with SIRIUS (v4.4).

### Quantification of flavonoids

Standards for naringenin chalcone (catalog number 33806, Cayman Chemicals), naringenin (N5893, Sigma Aldrich), and eriodictyol (catalog number 26803, Cayman Chemicals) were used to determine the content of these metabolites in frozen leaf tissue. Chloroform partitioned methanol-water extract was prepared from 100 mg of ground leaf tissue as described for the metabolome analysis, and 20 μL injected into UHPLC-MS/MS. The metabolites were quantified using an external calibration curve prepared using authentic standards.

### Pathway analysis

Accurate mass was used as input data for pathway enrichment analysis using the Functional Analysis module in MetaboAnalyst v6.0 (Li et al., 2013). Analysis was performed using negative mode, 10 ppm of mass tolerance, and a *p*-value cut-off of 0.05 for the *Mummichog* algorithm, and mapped to the pathway library of Arabidopsis from Kyoto Encyclopedia of Genes and Genomes.

### Identification of genes regulating metabolites and co-expression analysis

Genes encoding enzymes catalyzing the biosynthesis of metabolites were obtained from CornCyc 10.0.1 (https://www.plantcyc.org) online database. For metabolites associated with more than one gene, maize genes reported encoding these enzymes were identified from literature searches. Subsequently, only those genes that were expressed in the maize leaves (FPKM ≥ 2) during the post-flowering period in our expression dataset (Sekhon et al., 2019) were retained.

Co-expression networks were constructed using the WGCNA approach (Langfelder and Horvath, 2008) to analyze gene expression data from 43,868 genes across 112 senescence samples previously sequenced and submitted to SRA including PRJNA505891 (Sekhon et al., 2019) and PRJNA896566 (Kumar et al., 2023). To ensure robust and meaningful co-expression relationships, genes with FPKM = 0 in more than 30% of samples and FPKM values of <1 in all samples were filtered out. A soft-thresholding power of 6, chosen based on the scale-free topology criterion, was applied to create an adjacency matrix that captures the pairwise co-expression relationships between genes. The adjacency matrix was transformed into a topological overlap matrix (TOM) to account for shared neighbors and enhance the robustness of the network. Genes were hierarchically clustered based on the TOM, and co-expression modules were identified using the dynamic tree cut algorithm (Langfelder and Horvath, 2008) with a minimum module size of 30 genes. After network construction, genes with fewer than five co-expressing partners were removed. The filtered network and a node attribute file containing metadata on co-expressing partners were exported in edge list format for visualization and further analysis in Cytoscape (Shannon et al., 2003).

### Statistical analysis

Changepoint analysis was conducted to identify the timing at which each physiological parameter started to change and the rate of such change. This analysis was accomplished by fitting the following linear model for each inbred line separately:

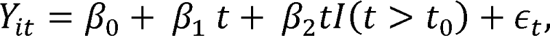

where Y_it_ is the physiological parameter (e.g., SPAD, Fv/Fm etc.) measured at time *t* on the *i*th replicate, β_0_ is the usual intercept, β_1_ and β_2_ are slope parameters, l(.) is the indicator function, and t_0_ is the possible changepoint. A few comments are warranted. First, the formulation of the changepoint model permits a slope of β_1_ in time for the physiological parameter up to time t_0_ and a slope of β_1_ + β_2_thereafter, thus capturing the rate of change attributable to senescence. Second, this model is fit for a grid of potential change point times, that is t_0_ ∈ [27, 27.1, …, 35.9, 36], and the optimal change point is selected via the configuration that maximizes R^2^ value. Third, we may identify the extent of the rate of change by examining the estimate of β_2_ with the values of this estimate equating to changes in the trend.

Marginal analysis was conducted to identify metabolites related to each of the physiological parameters. This analysis was accomplished by fitting the following linear model

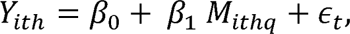

where Y_ith_ is the physiological parameter measured at time t on the ith replicate of the hth inbred line and M_ithq_ is the corresponding level of the qth metabolite. This model was fit considering all available metabolite levels to identify potential marginal associations between the metabolites and the various measures of senescence. Metabolome data was log-transformed to fit the assumption of normality for all analyses.

## SUPPLEMENTARY DATA

Supplementary Figure S1. Genetic diversity of inbred lines used in the study.

Supplementary Figure S2. Metabolic diversity in primary and secondary metabolome captured in the current study.

Supplementary Figure S3. Differentially abundant secondary metabolites in staygreen and non-staygreen inbred lines at different leaf development stages.

Supplementary Figure S4. Pathway enrichment analysis of secondary mass features concentrations at 33 days after anthesis in staygreen and non-staygreen inbred lines.

Supplementary Figure S5. Change in nitrogen content in staygreen and non-staygreen inbred lines at different leaf development stages during senescence

Supplementary Table S1: Marginal analysis of primary metabolites for different physiological parameters.

Supplementary Table S2: Compound validation of mass features and their p-values for different physiological phenotypes.

Supplementary Dataset S1. Phenotypic characterization of diverse inbred lines based on different physiological parameters.

Supplementary Dataset S2. Association of identified mass features with different physiological parameters.

Supplementary Dataset S3. Identification of genes encoding enzymes catalyzing the biosynthesis of different primary and secondary metabolites using CornCyc database.

## ACKNOWLEDGMENTS

We thank Rebecca Bishop, Laura Cicarelli, and many talented undergraduate students for assisting in preparing leaf tissues for metabolomic and C and N analyses. We would like to thank Elizabeth Leonard for her assistance with the metabolomic analyses.

## FUNDING

This work was supported in part by NSF grant #OIA-1826715 to RS.

## CONFLICT OF INTEREST STATEMENT

The authors declare no competing interests.

## AUTHOR CONTRIBUTION

R.S.S. designed the research; M.S.B. performed the research; M.S.B., B.K., R.K., N.T., and C.M. analyzed the data; and M.S.B. and R.S.S. wrote the article with help from all authors.

## Notes

### Competing Interest Statement

The authors have declared no competing interest.

