## Supplementary Materials for "Temporal analysis of physiological phenotypes identifies novel metabolic and genetic underpinnings of senescence in maize"

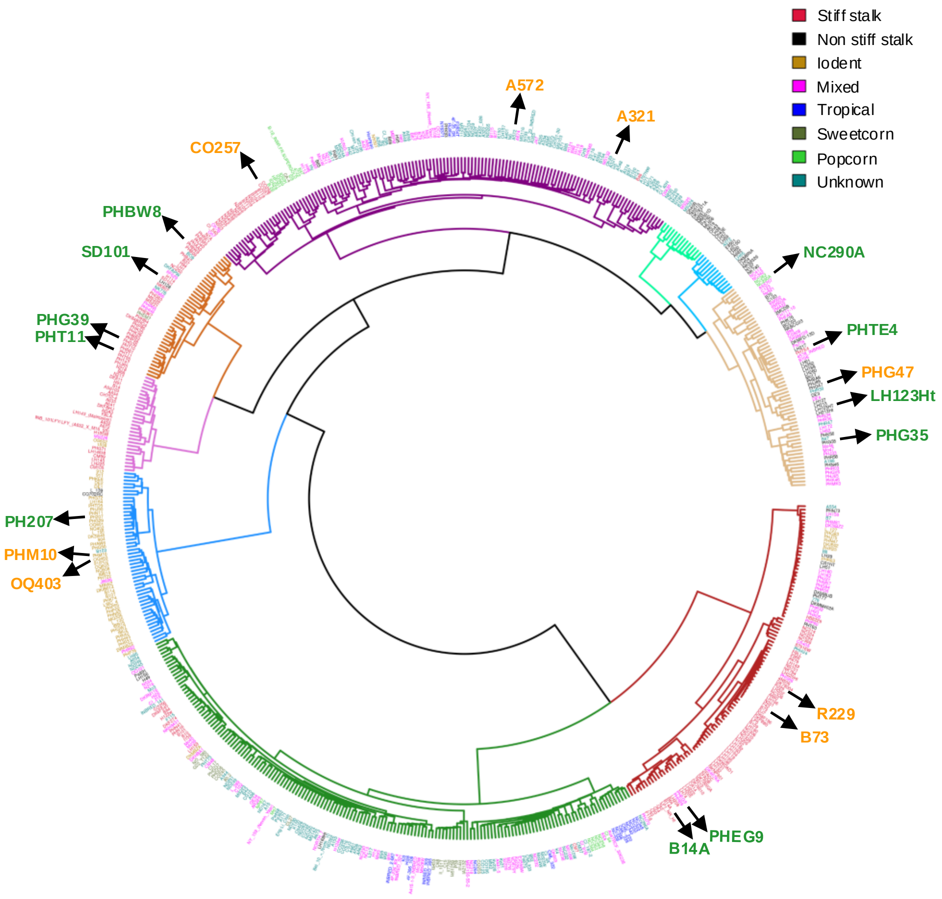


**Supplementary Figure S1:** Genetic diversity of inbred lines used in the study. Dendrogram of a subset of 603 inbred lines of the maize Wisconsin diversity panel showing the phylogenetic relationship between the inbreds. The entire subset is divided into 9 clusters as indicated by the colored branches and the inbred lines are classified into 8 germplasm groups as mentioned in the legend on the top right. The genetic grouping and location of the 19 inbred lines, included in the study, within the dendrogram are depicted by solid arrows and the staygreen and non-staygreen inbred lines are color coded in green and orange, respectively.

**A**

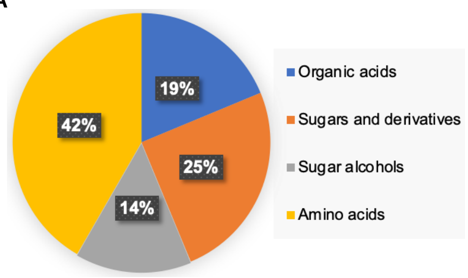


**B**

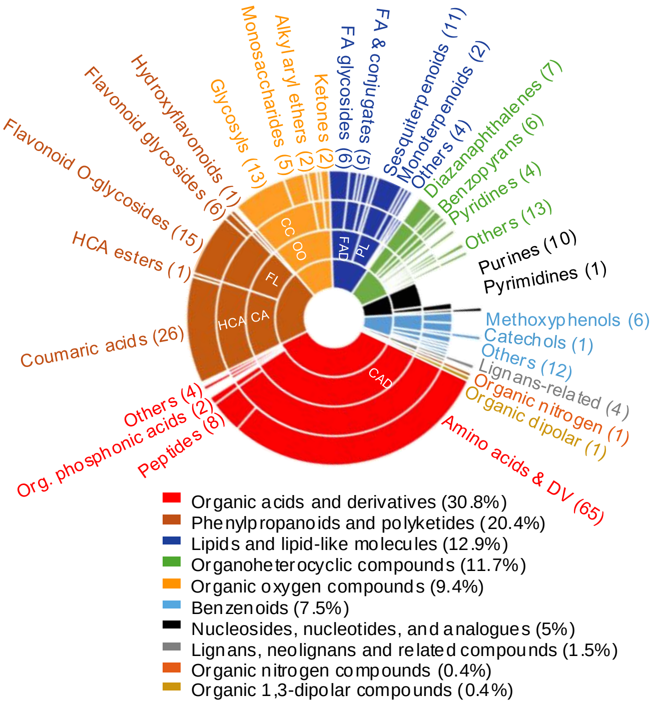


**Supplementary Figure S2**: Metabolic diversity in primary and secondary metabolome captured in the current study. A, Major primary metabolite classes. B, CANOPUS classification of secondary mass features annotated by the software. Each color represents a different chemical class. Moving away from the center, each layer denotes further sub-classification of a chemical class. Abbreviations: cinnamic acids, CA; carbohydrates and derivatives, CC; carboxylic acids and derivatives, CAD; derivatives, DV; fatty acid, FA; fatty acid derivatives, FAD; flavonoids, FL; Hydroxycinnamic acids, HCA; organooxygen compounds, OO; prenol lipids, PL.

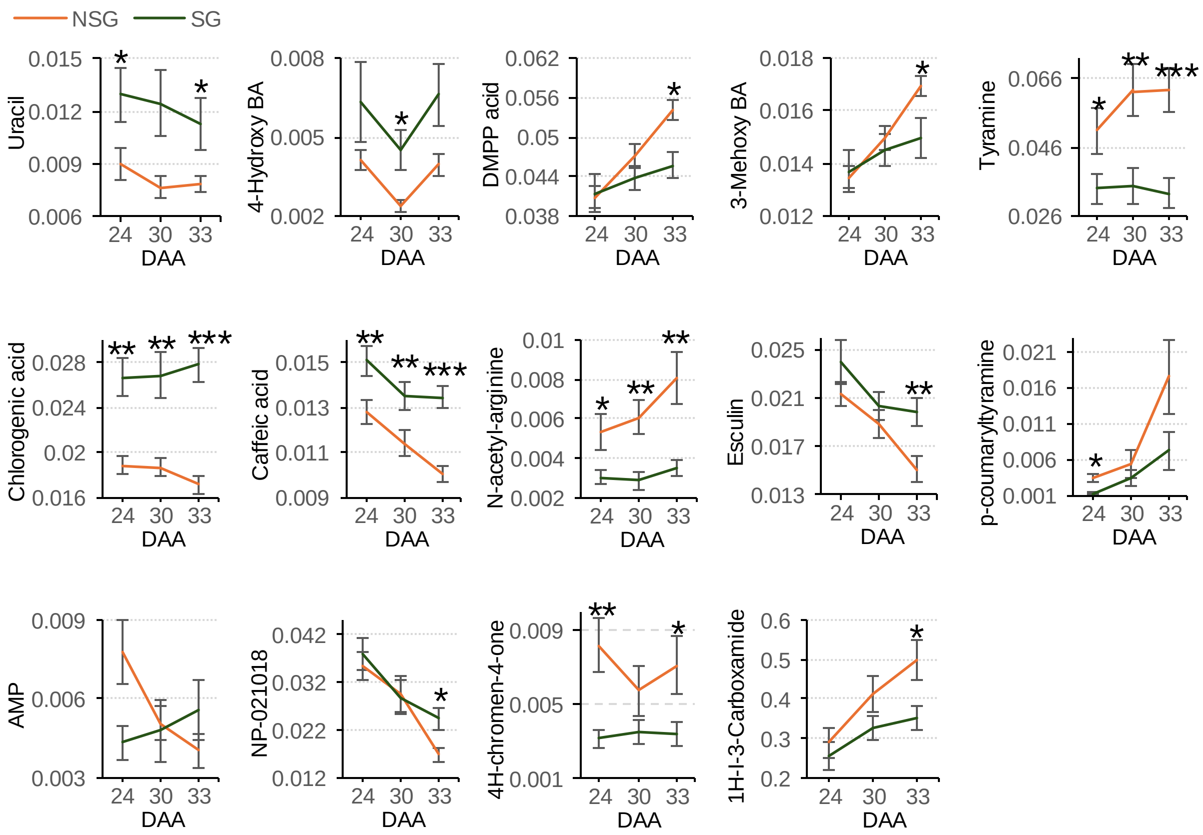


**Supplementary Figure S3**: Differentially abundant secondary metabolites in staygreen (darkgreen lines) and non-staygreen (orange lines) inbred lines at different leaf development stages. One, two, and three asterisks represent the student t-test significance level at $p \leq 0.05$, $p \leq 0.01$, and $p \leq 0.001$, respectively.

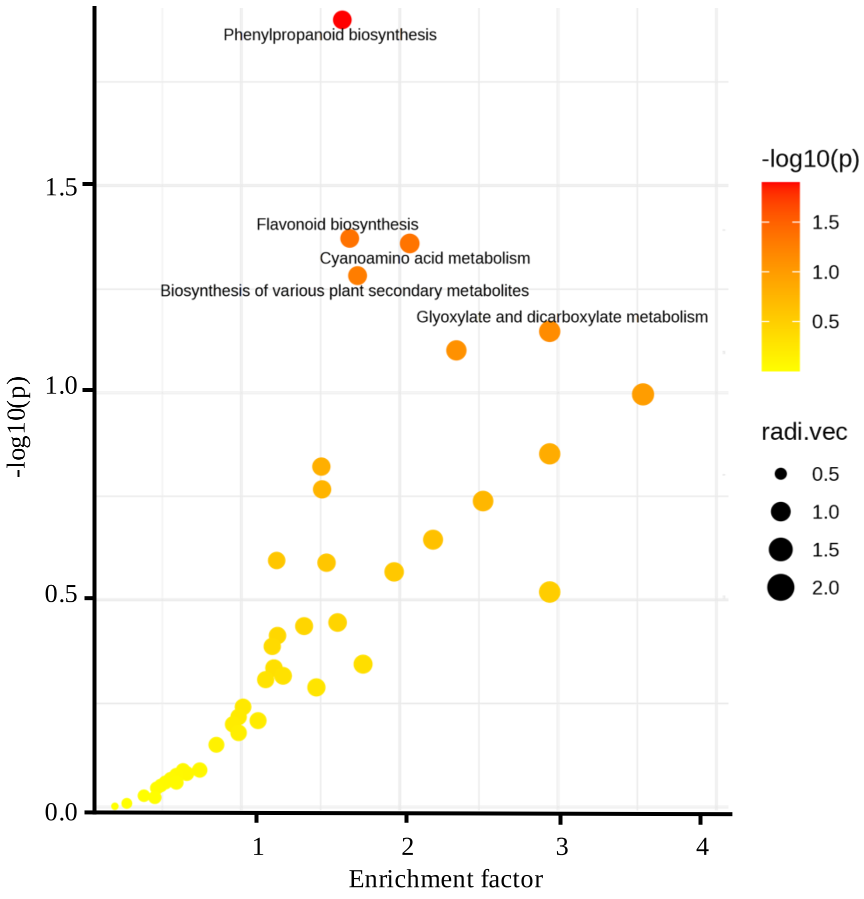


**Supplementary Figure S4**: Pathway enrichment analysis of secondary mass features concentrations at 33 days after anthesis in staygreen and non-staygreen inbred lines.

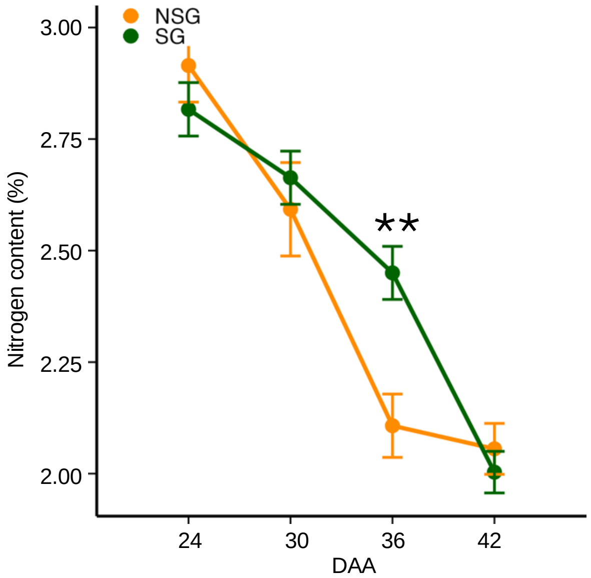


**Supplementary Figure S5**: Change in nitrogen content in staygreen (sg; darkgreen lines) and non-staygreen (nsg; orange lines) inbred lines at different leaf development stages during senescence. Two asterisks represent the student t-test level of significance at $p \leq 0.01$.

**Supplementary Table S1**: Marginal analysis of primary metabolites for different physiological parameters (p-value cut off from significant association 0.05)

| Metabolite | Class | p value (Fv/Fm | p value (SPAD | p value (CO2 assimilation | p value (ROS) | Significant (Fv/Fm | Significant (SPAD | Significant (CO2 assimilatio | Significant (ROS | DA |
|---|---|---|---|---|---|---|---|---|---|---|
| L-Isoleucine | Amino Acid | 1.21E-03 | 7.06E-09 | 2.85E-17 | 2.53E-03 | X | X | X | X | DA |
| L-Phenylalanine | Amino Acid | 2.98E-04 | 1.07E-16 | 1.05E-16 | 3.66E-02 | X | X | X | X | DA |
| L-Lysine | Amino Acid | 2.71E-04 | 3.68E-26 | 5.91E-25 | 4.41E-02 | X | X | X | X | DA |
| L-Arginine | Amino Acid | 3.54E-05 | 3.03E-14 | 3.16E-24 | 7.08E-02 | X | X | X | NS | DA |
| L-Leucine | Amino Acid | 4.61E-02 | 4.43E-08 | 8.08E-17 | 1.78E-01 | X | X | X | NS | DA |
| L-Methionine | Amino Acid | 9.01E-04 | 6.46E-12 | 4.80E-12 | 3.99E-03 | X | X | X | X | |
| Asparagine | Amino Acid | 1.11E-02 | 1.33E-13 | 2.75E-10 | 1.71E-03 | X | X | X | X | |
| L-Tyrosine | Amino Acid | 1.79E-02 | 1.68E-06 | 1.33E-04 | 7.20E-08 | X | X | X | X | |
| L-Valine | Amino Acid | 2.21E-02 | 4.72E-04 | 4.38E-07 | 1.86E-06 | X | X | X | X | |
| L-Aspartic-acid | Amino Acid | 1.35E-03 | 2.13E-04 | 3.67E-06 | 2.64E-02 | X | X | X | X | |
| L-Glutamine | Amino Acid | 7.87E-05 | 1.27E-06 | 2.75E-02 | 8.93E-04 | X | X | X | X | |
| Alanine | Amino Acid | 7.30E-04 | 6.55E-03 | 6.22E-05 | 4.46E-02 | X | X | X | X | |
| L-Histidine | Amino Acid | 1.48E-01 | 5.90E-07 | 7.95E-05 | 1.85E-03 | NS | X | X | X | |
| Glycine | Amino Acid | 4.01E-02 | 1.52E-01 | 2.30E-02 | 1.07E-01 | X | NS | X | NS | |
| L-Threonine | Amino Acid | 4.40E-01 | 2.29E-01 | 6.61E-02 | 1.63E-07 | NS | NS | NS | X | |
| Cysteine | Amino Acid | 2.72E-02 | 7.24E-01 | 5.40E-02 | 1.52E-02 | X | NS | NS | X | |
| Serine | Amino Acid | 4.82E-01 | 2.75E-01 | 7.16E-02 | 1.02E-04 | NS | NS | NS | X | |
| L-Proline | Amino Acid | 1.80E-01 | 6.19E-02 | 6.99E-03 | 6.75E-01 | NS | NS | X | NS | |
| L-Tryptophan | Amino Acid | 1.76E-01 | 8.19E-01 | 3.93E-01 | 4.65E-04 | NS | NS | NS | X | |
| a-Ketoglutarate | Organic acid | 2.86E-02 | 8.59E-09 | 9.56E-14 | 6.20E-04 | X | X | X | X | DA |
| Pyruvate | Organic acid | 2.22E-01 | 6.98E-01 | 5.08E-02 | 4.74E-01 | NS | NS | NS | NS | DA |
| Citric-acid | Organic acid | 4.70E-02 | 5.18E-11 | 4.33E-10 | 5.68E-04 | X | X | X | X | |
| Isocitric-acid | Organic acid | 5.07E-02 | 7.23E-11 | 5.96E-10 | 5.20E-04 | NS | X | X | X | |
| Succinic-acid | Organic acid | 5.99E-03 | 1.12E-13 | 5.26E-04 | 4.03E-01 | X | X | X | NS | |
| Malic-acid | Organic acid | 3.61E-01 | 5.73E-02 | 1.38E-02 | 2.65E-10 | NS | NS | X | X | |
| Shikimic-acid | Organic acid | 6.64E-01 | 2.44E-02 | 9.61E-04 | 6.07E-03 | NS | X | X | X | |
| Fumaric-acid | Organic acid | 9.21E-01 | 1.48E-02 | 9.51E-01 | 1.76E-02 | NS | X | NS | X | |
| L-Glutamic acid | Organic acid | 5.29E-01 | 6.41E-02 | 1.20E-01 | 1.57E-01 | NS | NS | NS | NS | |
| Ascorbic acid | Organic acid | 1.92E-01 | 6.78E-01 | 2.98E-01 | 6.71E-02 | NS | NS | NS | NS | |
| Lactose | Sugar | 1.17E-02 | 1.17E-01 | 5.98E-04 | 3.55E-02 | X | NS | X | X | DA |
| Ribose | Sugar | 2.36E-01 | 1.57E-04 | 5.20E-05 | 2.69E-01 | NS | X | X | NS | DA |
| Sucrose | Sugar | 8.44E-01 | 4.13E-01 | 8.56E-01 | 4.51E-01 | NS | NS | NS | NS | DA |
| D-Maltose | Sugar | 4.27E-19 | 5.22E-29 | 7.03E-32 | 4.45E-06 | X | X | X | X | |
| Arabinose | Sugar | 1.21E-04 | 1.47E-12 | 7.52E-11 | 3.13E-05 | X | X | X | X | |
| Glucose-6-Phosphate | Sugar | 1.52E-01 | 2.95E-07 | 6.39E-02 | 4.46E-02 | NS | X | NS | X | |
| Fructose | Sugar | 2.74E-01 | 4.36E-02 | 1.18E-02 | 3.91E-02 | NS | X | X | X | |
| Xylose | Sugar | 8.34E-05 | 2.14E-28 | 3.35E-17 | 4.21E-01 | X | X | X | NS | |
| Fructose-6-Phosphate | Sugar | 5.70E-04 | 8.81E-09 | 6.96E-08 | 6.23E-01 | X | X | X | NS | |
| Glucose | Sugar | 5.30E-01 | 2.69E-01 | 1.47E-01 | 4.66E-02 | NS | NS | NS | X | |
| Mannose | Sugar | 9.40E-01 | 5.00E-01 | 8.54E-02 | 3.13E-01 | NS | NS | NS | NS | |
| Galactose | Sugar | 9.86E-01 | 5.49E-01 | 6.28E-01 | 2.71E-01 | NS | NS | NS | NS | |
| Iso-erythritol | Sugar Alchohol | 1.15E-14 | 1.40E-20 | 3.69E-22 | 3.75E-12 | X | X | X | X | DA |
| Arabitol | Sugar Alchohol | 1.01E-11 | 2.77E-22 | 2.05E-37 | 8.65E-06 | X | X | X | X | DA |
| Mannitol | Sugar Alchohol | 2.68E-22 | 8.97E-40 | 2.19E-51 | 3.58E-05 | X | X | X | X | DA |
| Galactitol | Sugar Alchohol | 8.03E-05 | 5.71E-07 | 4.52E-24 | 3.89E-02 | X | X | X | X | DA |
| Sorbitol | Sugar Alchohol | 1.01E-03 | 2.90E-06 | 5.28E-04 | 2.38E-03 | X | X | X | X | |
| Maltitol | Sugar Alchohol | 3.98E-02 | 1.15E-01 | 2.70E-01 | 8.74E-01 | X | NS | NS | NS | |
| Xylitol | Sugar Alchohol | 2.87E-01 | 3.29E-03 | 9.18E-01 | 8.60E-01 | NS | X | NS | NS | |

X and NS denotes significant and non-significant association with physiological phenotype, respectively

DA represents differential abundance of metabolite between the groups of staygreen and non-staygreen inbred lines

**Supplementary Table S2**: Compound validation of mass features and their p-values for different physiological phenotypes (X denotes significantly associated)

| Class | Mass feature | Metabolite Validation | Category | p_value_Fv/Fm | p_value_SPAD | p_value_CO2 assimil. | p_value_ROS | DA | Significant (Fv/Fm) | Significant (SPAD) | Significant (CO2 assimil.) | Significant (ROS) |
|---|---|---|---|---|---|---|---|---|---|---|---|---|
| Phenylpropanopoids | 323.139_10.32 | N-(p-Coumaroyl) serotonin | Amine | 7.47E-10 | 7.00E-38 | 1.83E-26 | 1.53E-03 | - | X | X | X | X |
| Other | 144.08_7.52 | 2-Naphthylamine | Amine | 1.97E-12 | 2.09E-21 | 1.37E-35 | 1.05E-02 | Yes | X | X | X | X |
| Phenylpropanopoids | 284.128_10.49 | (2E)-3-(4-Hydroxyphenyl)-N-[2-(4-hydroxyphenyl)ethyl]acrylamide (p-coumaryl tyramine) | Amine | 4.56E-06 | 4.63E-22 | 1.50E-26 | 1.52E-03 | Yes | X | X | X | X |
| Phenylpropanopoids | 284.128_10.3 | (2E)-3-(4-Hydroxyphenyl)-N-[2-(4-hydroxyphenyl)ethyl]acrylamide (p-coumaryl tyramine) | Amine | 7.26E-04 | 2.64E-17 | 5.30E-27 | 1.14E-04 | Yes | X | X | X | X |
| Benzaldehydes | 137.059_8.68 | 3-Methoxybenzaldehyde | Benzaldehydes | 3.53E-02 | 3.19E-22 | 9.28E-22 | 3.40E-01 | Yes | X | X | X | - |
| Benzaldehydes | 123.044_2.8 | 4-Hydroxybenzaldehyde | Benzaldehydes | 8.78E-01 | 2.03E-01 | 9.06E-03 | 7.68E-01 | Yes | - | - | X | - |
| Phenylpropanopoids | 341.086_7.21 | Esculin | Coumarins | 1.89E-03 | 9.49E-14 | 2.42E-20 | 7.99E-02 | Yes | X | X | X | - |
| Phenylpropanopoids | 449.108_9.6 | Cynaroside (luteolin-7-O-glucoside) | Flavone | 2.59E-01 | 1.61E-01 | 9.58E-01 | 8.32E-06 | - | - | - | - | X |
| Phenylpropanopoids | 273.075_11.54 | Naringeninchalcone | Flavonoid | 7.66E-09 | 1.50E-26 | 1.12E-17 | 2.66E-01 | - | X | X | X | - |
| Phenylpropanopoids | 611.161_8.53 | Rutin | Flavonoid | 1.65E-06 | 1.13E-08 | 2.29E-05 | 3.79E-06 | - | X | X | X | X |
| Phenylpropanopoids | 287.054_8.66 | Kaempferol, In-source fragment of Kaempferol-O-deoxyhexose glucoside (m/z 595.1661) | Flavonoid | 2.47E-02 | 6.12E-08 | 1.36E-05 | 1.97E-06 | - | X | X | X | X |
| Phenylpropanopoids | 289.07_9.59 | Eriodictyol | Flavonoid | 2.12E-02 | 2.34E-17 | 5.20E-14 | 8.99E-01 | Yes | X | X | X | - |
| Phenylpropanopoids | 355.102_6.94 | Chlorogenic acid | Hydroxycinnamates | 2.82E-02 | 1.16E-22 | 4.13E-40 | 1.88E-01 | Yes | X | X | X | - |
| Phenylpropanopoids | 163.039_8.1 | Caffeic acid, water loss | Hydroxycinnamates | 4.73E-03 | 4.31E-25 | 6.67E-21 | 1.17E-01 | - | X | X | X | - |
| Phenylpropanopoids | 181.049_6.94 | Chlorogenic acid (caffeic acid in-source fragment) | Hydroxycinnamates | 1.70E-02 | 1.11E-18 | 1.35E-26 | 4.94E-02 | Yes | X | X | X | X |
| Phenylpropanopoids | 163.039_6.94 | Chlorogenic acid (caffeic acid water loss in-source fragment) | Hydroxycinnamates | 4.40E-02 | 1.99E-20 | 1.10E-35 | 1.57E-01 | Yes | X | X | X | - |
| Phenylpropanopoids | 195.065_7.81 | Ferulic acid | Hydroxycinnamates | 6.42E-05 | 8.16E-01 | 4.16E-03 | 2.33E-11 | - | X | - | X | X |
| Phenylpropanopoids | 195.065_9.01 | Ferulic acid | Hydroxycinnamates | 7.80E-01 | 6.55E-22 | 1.82E-13 | 9.59E-04 | - | - | X | X | X |
| Phenylpropanopoids | 225.075_8.05 | Sinapinic acid | Hydroxycinnamates | 6.26E-02 | 5.31E-01 | 9.42E-02 | 9.11E-07 | - | - | - | - | X |
| Phenylpropanopoids | 325.092_7.82 | Caffeoyl-O-glucoside | Hydroxycinnamates | 7.54E-02 | 1.21E-01 | 1.65E-01 | 2.55E-06 | - | - | - | - | X |
| Phenylpropanopoids | 355.102_6.01 | Chlorogenic acid | Hydroxycinnamates | 1.80E-01 | 2.86E-05 | 4.23E-09 | 1.04E-01 | Yes | - | X | X | - |
| Phenylpropanopoids | 225.075_8.48 | Sinapinic acid | Hydroxycinnamates | 2.87E-01 | 2.67E-01 | 2.41E-01 | 1.89E-07 | - | - | - | - | X |
| Phenylpropanopoids | 225.075_7.43 | Sinapinic acid derivative, In-source fragment [M-179.0796+H]+ of m/z 404.1553 (identity of parent not confirmed) | Hydroxycinnamates | 1.05E-01 | 9.60E-01 | 7.63E-01 | 4.35E-07 | - | - | - | - | X |
| Other | 206.044_6.58 | Xanthurenic acid | Kynurenines | 3.50E-02 | 6.55E-03 | 4.51E-01 | 2.47E-09 | - | X | X | - | X |
| Other | 190.049_7.37 | Kynurenic acid | Kynurenines | 1.38E-01 | 1.92E-04 | 2.25E-01 | 8.53E-09 | - | - | X | - | X |
| Nucleotide | 113.034_2.97 | Uracil | Nucleotide | 4.22E-03 | 5.60E-16 | 1.84E-11 | 9.77E-02 | Yes | X | X | X | - |
| Other | 227.127_9.72 | NP-021018 | Other | 1.31E-11 | 2.13E-41 | 1.55E-28 | 3.16E-01 | Yes | X | X | X | - |
| Other | 160.075_4.73 | 1H-indene-3-carboxamide | Other | 3.81E-06 | 6.82E-36 | 3.54E-30 | 6.46E-01 | Yes | X | X | X | - |
| Other | 227.127_9.57 | NP-021018 | Other | 5.03E-10 | 5.56E-27 | 3.85E-17 | 2.26E-01 | - | X | X | X | - |
| Other | 227.127_7.9 | NP-021018 | Other | 2.63E-05 | 3.15E-09 | 6.00E-19 | 1.53E-06 | Yes | X | X | X | X |
| Other | 123.08_8.68 | dimethoxyphenylpropenoic acid | Other | 6.00E-03 | 5.24E-24 | 3.53E-26 | 2.29E-01 | Yes | X | X | X | - |
| Other | 382.171_6.37 | Pantothenic acid 4'-O-b-D-glucoside | Other | 1.10E-03 | 9.08E-02 | 3.11E-02 | 1.02E-06 | - | X | - | X | X |
| Other | 285.169_9.02 | NP-016437 | Other | 1.96E-09 | 2.61E-17 | 5.18E-07 | 6.24E-01 | - | X | X | X | - |
| Other | 217.129_2.78 | N-α-L-Acetyl-arginine | Other | 2.73E-02 | 1.04E-10 | 2.31E-17 | 4.88E-01 | Yes | X | X | X | - |
| Other | 138.091_3.35 | Tyramine | Other | 6.45E-01 | 5.84E-09 | 2.73E-09 | 2.47E-04 | Yes | - | X | X | X |
| Other | 369.118_6.46 | (3R,5R)-1,3,5-Trihydroxy-4-{[(2E)-3-(4-hydroxy-3-methoxyphenyl)-2-propenoyl]oxy}cyclohexanecarboxylic acid | Other | 2.38E-01 | 9.05E-02 | 5.62E-01 | 1.45E-09 | - | - | - | - | X |
| Other | 387.201_8.56 | (4S)-4-hydroxy-3,5,5-trimethyl-4-[(1E)-3-{[(2R,3R,4S,5S,6R)-3,4,5-trihydroxy-6-(hydroxymethyl)oxan-2-yl]oxy}but-1-en-1-yl]cyclohex-2-en-1-one | Other | 2.46E-01 | 4.50E-01 | 4.26E-01 | 7.16E-06 | - | - | - | - | X |
| Other | 463.123_9.05 | 5,7-dihydroxy-6-methoxy-2-(4-{[(2S,3R,4S,5S,6R)-3,4,5-trihydroxy-6-(hydroxymethyl)oxan-2-yl]oxy}phenyl)-4H-chromen-4-one | Other | 7.12E-02 | 7.91E-01 | 3.32E-01 | 3.17E-02 | Yes | - | - | - | X |
| Other | 195.137_10.26 | Sedanolide derivative, In-source fragment of m/z 241.1434 | Other | 2.77E-01 | 1.99E-06 | 7.59E-02 | 4.89E-01 | - | - | X | - | - |
| Other | 144.101_2.76 | 1-Aminocyclohexanecarboxylic acid | Other | 3.31E-01 | 2.55E-01 | 4.10E-01 | 7.45E-02 | - | - | - | - | - |
| Other | 348.07_2.77 | Adenosine 5'-monophosphate | Other | 7.26E-01 | 7.22E-01 | 6.25E-02 | 8.51E-01 | Yes | - | - | - | - |
| Terpene | 153.127_8.55 | Pulegone | Terpene | 5.41E-01 | 8.69E-01 | 7.73E-06 | 4.97E-01 | - | - | - | X | - |
